# A Linear Mixed Effects Model for Evaluating Synthetic Gene Circuits

**DOI:** 10.1101/2024.12.30.630778

**Authors:** Gina Partipilo, Sarah M. Coleman, Yang Gao, Ismar E. Miniel Mahfoud, Claus O. Wilke, Hal S. Alper, Benjamin K. Keitz

**Author notes:** Denotes co-contributions; **Corresponding Author:** Hal S. Alper, Benjamin K. Keitz, **Email:**. Denotes co-contributions (Gina Partipilo, Sarah M. Coleman). **Author Contributions:** GP, SMC, HSA, and BKK conceived the project. GP and SMC performed data scrape, analysis, and conceptualized. SMC performed statistical analysis, data simulation, and mixed effects modeling. GP cloned nested repressor gate constructs and characterized performance. YG provided electrical engineering expertise and perspective. IEMM created and designed the online interface. COW provided expertise on statistical analysis. GP and SMC wrote the manuscript with input from all the authors. **Competing Interest Statement:** There are no competing interests.

## Abstract

A significant advancement in synthetic biology is the development of synthetic gene circuits with predictive Boolean logic. However, there is no universally accepted or applied statistical test to analyze the performance of these circuits. Many basic statistical tests fail to capture the predicted logic (OR, AND, etc.) and most studies neglect statistical analysis entirely. As synthetic gene circuits shift toward advanced applications, primarily in computing, biosensing, and human health, it is critical to standardize the statistical methods used to evaluate gate success. Here, we propose the application of a linear mixed effects model to analyze and quantify genetic Boolean logic gate performance. First, we analyzed 144 currently published Boolean logic gates for trends and used unsupervised machine learning (k-means clustering) to validate the statistical model. Next, we utilized the model to generate estimates for the fixed effect of the ON state, ß, as a general descriptor of the Boolean nature of a circuit and used Monte Carlo simulations to recommend sample sizes for evaluating gate performance. Finally, we examined ß as a holistic metric for circuit performance using a series of nested repressor OR gates with intentionally degraded performance. We observed a linear correlation between ß and the predicted translation rate, highlighting the use of ß for the forward design of new Boolean gates. In summary, we utilized a linear mixed effects model to describe synthetic gene circuits and determined that the fixed effect, ß, is an appropriate descriptor of gate behavior that can be used to statistically evaluate performance.

**Significance statement:** There is no standard method for statistically evaluating the success of biological Boolean logic gates and common statistical tests are not used or incorrectly applied. We propose the use of a linear mixed effects model to estimate the value for ß, a parameter which describes the difference between ON and OFF gate states. A null hypothesis of ß = 0 can then be tested to determine if there is a statistically significant difference between the two states.

## Introduction

2-input Boolean logic gates utilize a set of input signals (1s and 0s) to produce an output signal or state (1 or 0)^1^. Boolean algebra has been applied for the design of digital circuits, where the inputs take the form of high (1) and low (0) voltage or the presence or absence of current in a given gate^2,3^. When connected in circuits, logic gates can combine to perform advanced computation. As a result, semiconductor progress and circuit design have been driven by the need for component miniaturization, faster switching speed, higher power efficiency, minimization of noise, and increased reliability^4^. Replicating these advances in living systems, synthetic biology has developed genetic circuits and logic gates, akin to electrical circuits, which transduce chemical, biological, or optical signals into controlled biological outputs, typically fluorescence. Specifically, native and engineered single-input gates can form inducible gene expression networks^5,6^, which have been extrapolated and engineered to created multi-input synthetic Boolean circuits. Synthetic gene circuits can be controlled by DNA ^7–10^, including quorum sensing^11–14^, recombinases^15,16^, and transcription factors^17–22^; RNA^23–26^; and proteins ^27–35^; and have been deployed across multiple phyla including bacteria^11,12,23,24,36–39^, mammalian cells^25,27,40–42^, and plants^17^.

Although synthetic gene circuits seek to mimic electrical circuits, they differ in several critical ways (Figure 1). For electrical circuits, voltage and current are standardized and absolute units of measurement. The response time of electrical circuits is often fast (micro-to nanoseconds), and they typically contain no state memory. As such, individual transistors within the circuit are assumed to operate identically and independently. Conversely, biological circuits operate over much longer time scales (minutes to hours). Because circuits may function in different parts of the cell cycle, they may also induce permanent or semi-permanent shifts in the biological state, which endows the system with an inherent state memory. Furthermore, the characterization of individual components within synthetic gene circuits (*i*.*e*., promoter/repressor^6,32^, translational repression^24^, etc.) is not necessarily representative of their behavior in a higher-order gate due to biological cross-talk and interdependence of the two signals. Finally, the output flexibility of genetic Boolean logic gates yields complex phenotypes that are difficult to standardize, making the output more variable and indirect compared to current or voltage. These factors make forward gate design, optimization, and troubleshooting more challenging compared to electrical systems. Nonetheless, many synthetic gene circuits have been developed with a fluorescence or absorbance^43^ as an output, and have been applied to control cell morphology^35,39^ and as cancer therapeutics^44^.

**Figure 1.**
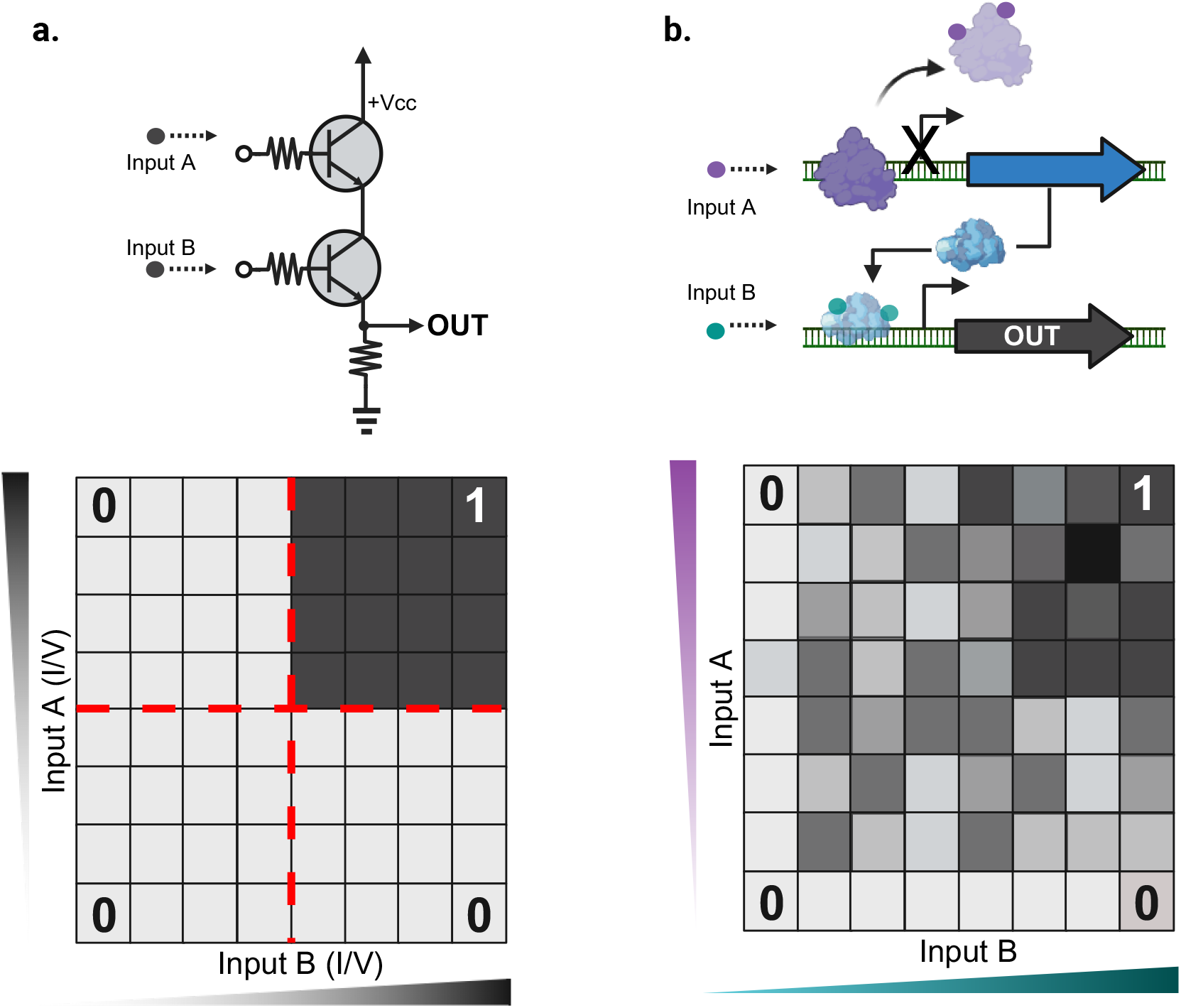
Boolean logic in biology versus electrical systems of an AND gate. **a**. Schematic describes the inputs (voltage or current) of a transistor-based Boolean logic gate and the tunable nodes with potential interactions before giving an output (voltage or current). **b**. Schematic describes the complex and flexible inputs of a biological Boolean logic gates and the potential cross-talk between component yielding multiple different potential outputs. Bottom heat maps represent output variability.

While success for relatively simple gates may be intuitive, as the field increases in complexity of both regulation^45^ and application^44^, there will be an increased need for more quantitative success metrics—particularly when moving towards applications in human health. Here, we propose a linear-mixed effects model to quantify the performance of synthetic gene circuits. The parameters estimated by the model can be taken as individual descriptors of gate behavior (Figure 2), which can then be evaluated by a general linear hypothesis test for examining the fixed effect of state. We utilize this model to examine state-of-the-art genetic logic gates and determine that there is general agreement between well-performing gates and gates with fixed effects that are statistically greater than zero. To help guide experimental design, we propose sample sizes for well-performing gates through a series of Monte Carlo simulations. We implement these findings to provide an online resource which aids researchers in the analysis of their genetic Boolean gates. Users can input data into a table, the linear mixed effects model is completed and statistically evaluated, and the outputs plotted. Finally, we motivate the model’s use through the generation of a small panel of nested repressor gates. Ultimately, the model for evaluating gate success can be combined with other synthetic biology tools (RBS calculators^46^, computational Boolean Logic predictors^47^, and machine learning models) to help standardize and shape new genetic Boolean gate development.

**Figure 2.**
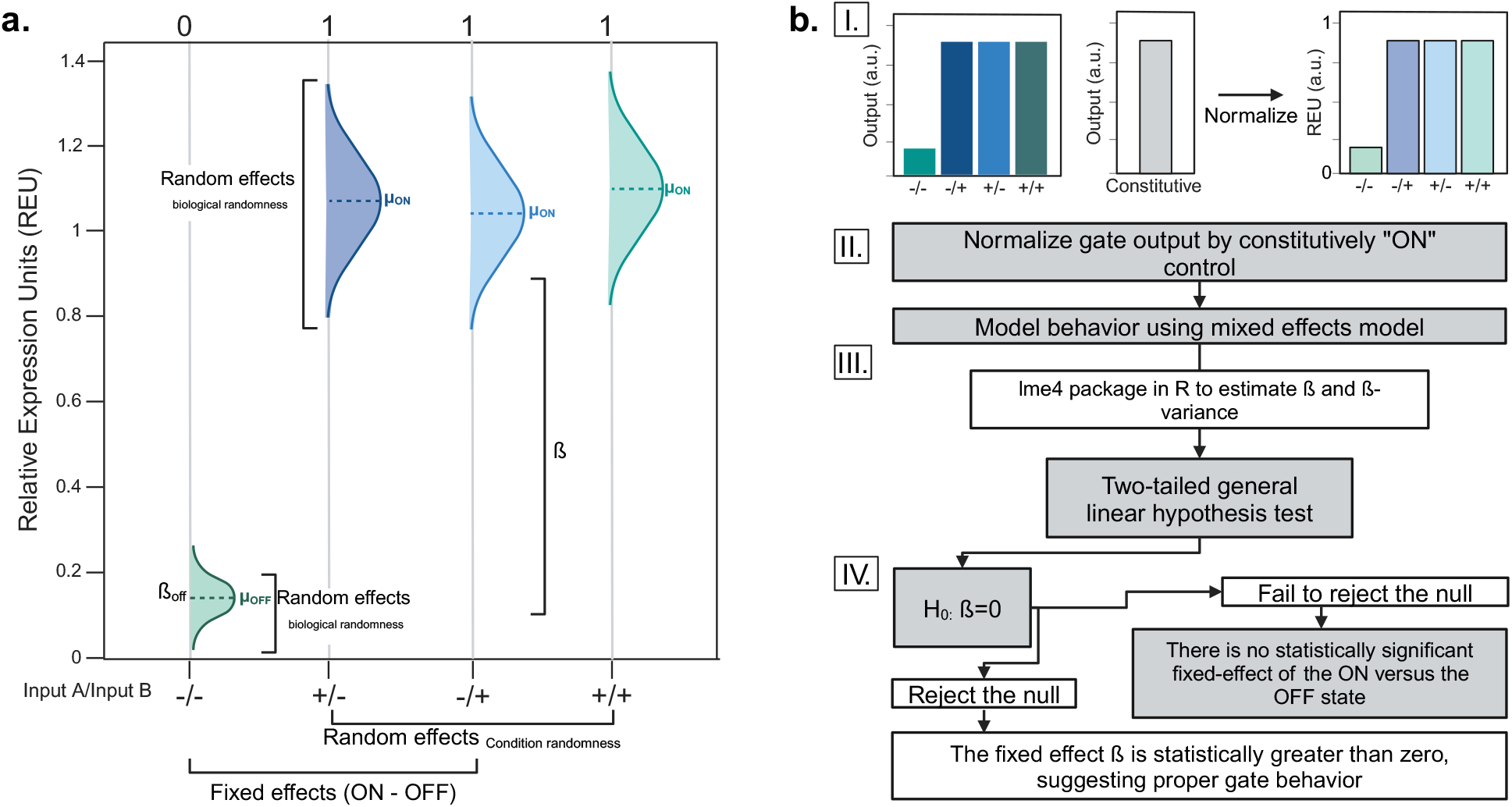
Description of a linear mixed effects model when applied to Boolean Logic gates. **a**. Given two inputs (A or B), the schematic describes outputs available in two different states (OFF ~ 0.1 REU, and ON ~1 REU). The gate described is an “OR” gate where a higher output is achieved in the presence of either input (A or B). The mixed effects model describes the combination of fixed and random effects for a Boolean logic gate. The fixed effect (ß) is interpreted as the amount of increase in signal between the OFF and ON states. **b**. The workflow describes a process for pre-treating raw Boolean logic data before fitting a linear mixed effects model to estimate ß and performing a hypothesis test to determine if ß is statistically greater than zero.

## Results

### A linear mixed effects model for modeling Boolean gate performance

The existing statistical tests used in the literature for evaluating synthetic gene circuits are primarily Student’s t-test and ANOVA, but even these are rarely applied. Unfortunately, these comparative statistics fail to quantify both the nature and performance of a circuit. We posit that a linear mixed effects model can more accurately describe gate behavior in a manner that goes beyond simply comparing the statistical significance of a singular ON/OFF state. In such a model, the “fixed” effect corresponds to output state (ON or OFF) while the “random” effect corresponds to variations between input states with the same output state. Linear-mixed effects models are common in biology^48^, particularly ecology^49–51^, and have also been applied to biomedical^52^ and genomic^53,54^ datasets. However, to the best of our knowledge they have not been used for evaluating Boolean logic gates. A significant advantage of using a linear mixed effects model is the ability to estimate a singular parameter (the fixed effect of ON over OFF, ß), which can be used to quantitatively compare different circuits. Following conventional notation in the field^55^, a linear mixed-effects model for genetic logic gate data can be described by the following^56^:

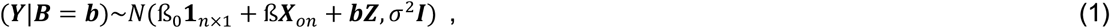

where the response vector ***Y*** within an input group (component of random effects variable ***B*** fixed at ***b***) is normally distributed around the sum of the fixed effect predictors ß_0_**1**_*n*×1_ and ß***X***_*on*_, and the random effects predictors ***bZ***. The final error term for the response vector ***Y*** is modeled as *σ*^2^***I***, where ***I*** represents the *n* _×_ *n* identity matrix. In this model formulation, **1**_*n*×1_ and ***X***_*on*_ are fragmentations of the fixed effects design matrix, with ***X***_*on*_ serving as an indicator variable (either 0 or 1) representing gate output, while ***Z*** represents the random effects design matrix. Each random effect may have different variances (see Materials and Methods). In this formulation, ß_0_ and ß represent the intercept (OFF state) and gate response (ON - OFF), respectively. The parameters estimated by the linear mixed effects model are all useful descriptors of circuit performance; however, as illustrated in Figure 2, we propose ß (the fixed effect of ON-OFF) as a single descriptor of overall gate behavior.

A ß estimate that is statistically different than zero (equivalent to rejecting the null hypothesis, H_0_: ß=0) indicates there is a difference between the OFF and ON populations. Broadly speaking, maximizing ß maximizes the differences between the OFF and ON states and is an appropriate metric to benchmark gate response. The results of this test are contextual, based on the mode of measurement, and it is critical to control for the random effect of input state. For example, fluorescence values which are dependent on the instrument used are often represented by very high arbitrary units. These values are much more likely to yield high ß estimation, due to the arbitrarily high units, making it difficult to compare between analogous genetic circuit systems. To circumvent this issue, data should be normalized in a way that benchmarks responses to reasonable values, and we suggest normalizing data with a positive control that lacks regulation to represent the highest “ON” level achievable^22,34,45,57^. In this case, ß estimates closer to 1 indicate that the system is behaving logically and displaying binary, multi-state behavior. However, in many application spaces it is difficult to achieve an appropriate positive control. In this case, ß estimates statistically greater than zero are still meaningful for indicating logical response but caution should be applied when deciding whether they indicate a truly binary response.

#### Box 1. Definitions of key parameters

The **fixed effect, ß**, quantifies the difference in response between ON and OFF states

The null hypothesis, **H**_**0**_**: ß=0**, is that the fixed effect of ON is equal to OFF

The ***p*-value** is the probability of observing these (or more extreme) data under the null hypothesis, ß=0.

### Conceptualizing the Biological Boolean logic gate space via unsupervised machine learning clustering

To demonstrate the utility of ß estimations as an appropriate statistical metric of success, we performed a meta-analysis of 144 gates published across 22 articles and evaluated the mean output for each input condition for multiple different gate types (OR, NOR, AND, NAND, IMPLY, NIMPLY, XOR, and XNOR). During this analysis, we found that many papers either commented on performance without performing statistical tests, or, in some cases, defined an arbitrary ON/OFF threshold to interpret results. The inputs and output conditions for each gate type (1s and 0s) are described in Supplementary Table S1. Due to the variety of output data (fluorescence, absorbance, band size, etc.), we first applied a normalization metric to the data (Figure 3a). The data were normalized by the average of the “ON” conditions, such that the average of the ON was equivalent to 1 and the OFF gates were less than 1 (Figure S2). This enabled a facile comparison between drastically different types of gate outputs and controlled for instrument-to-instrument variability. While we initially intended to scrape individual data observations instead of mean outputs, we found that the mean response with standard deviation was by far the most the most common data representation published in the literature.

**Figure 3.**
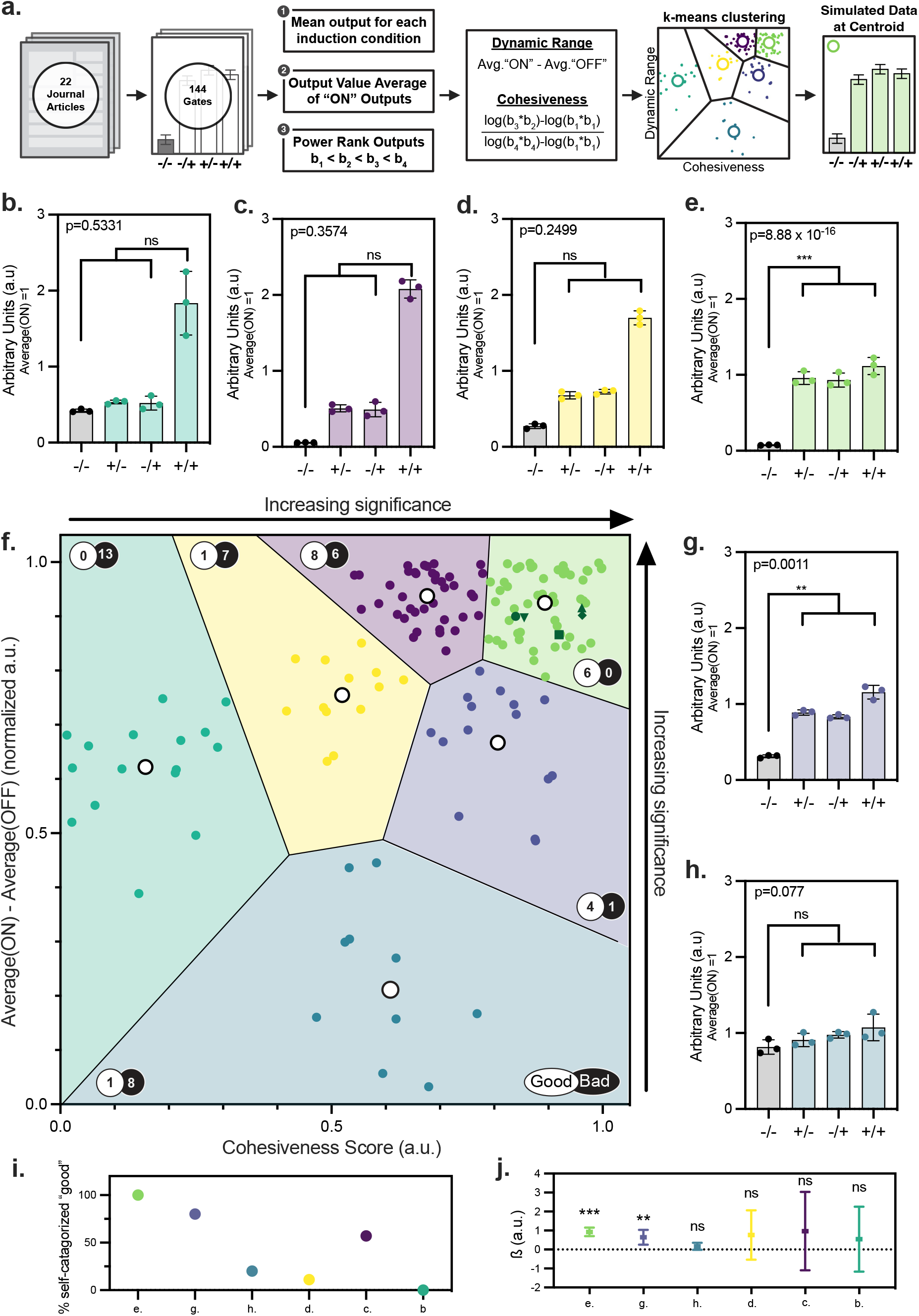
The fixed effects parameter ß holistically describes gate performance. **a**. Schematic dictating the workflow of the data scrape from 22 papers and 144 logic gates. **b.-e**. Simulated data for each of the centroid points in **f**. where the average of the “ON” values are normalized to 1. The variance for each data simulation is simulated as 10% of the mean (assuming a Gaussian distribution) and the results of the hypothesis test of ß after model fit are represented with p-values on each graph. Only the estimate of ß in **e**. is statistically different than zero at a significance level of 0.05. **f**. A two-dimensional description of gate performance for gates scraped to map the current output landscape of biological Boolean logic gates. Each gate is overlaid with a ratio of data that was self-categorized by the original authors of the scraped paper as being “good” to “bad”. The dark green data points indicate commercial CMOS Boolean Logic gates. The same data is reprinted in Figure S1. and overlaid with type of regulatory mechanism and gate type. Regions are plotted with Voronoi lines separating clusters and are the result of unsupervised machine learning. **g.-h**. Simulated data for each of the centroid points in **f**. where the average of the “ON” values is normalized to 1. The variance for each data simulation is 10% of the mean (assuming a Gaussian distribution) and the results of the hypothesis test of ß after model fit are represented with p-values on each graph. Only the estimate of ß in **g**. is statistically different than zero at a significance level of 0.05. **i**. Percentage of self-categorized “good” gates per Voronoi region. **j**. ß estimates resulting from model fit to simulated data for each of the centroid points (**b.-e**., **g**., **h**.) as a single descriptor of gate performance for each centroid. Error bars represent the 95% Confidence Interval (CI) for each ß estimate.

With the caveat that each gate contains an internal normalization to the average of the ON conditions, two qualifying descriptors were extracted from each gate. The first was the average of the output responses predicted to be ON (equivalent to 1 due to our normalization) minus the average of the output responses predicted to be OFF. An ideal gate, regardless of specific logic, would approach 1, reflecting the dynamic range achieved by the logical regulation. The second descriptor represents the spread and variability of the response outputs. Adapting a quantification of OR and AND gates proposed by Cox^58^ to evaluate the cohesiveness (logic) of a gate, we rank-ordered (smallest to largest, b_1_-b_4_) the mean response for each gate and used the following equations to determine the cohesiveness of each depending on the predicted gate function. Equation 2 was used for NAND, OR, and IMPLY, where there is only one OFF state; equation 3 for NOR, AND, and NIMPLY, where there is only one ON state; and equation 4 for XOR and XNOR, where there are two ON and OFF states.

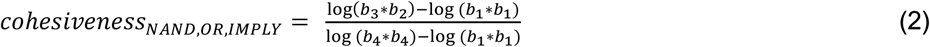

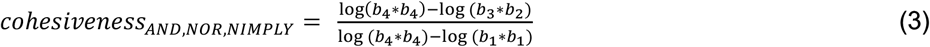

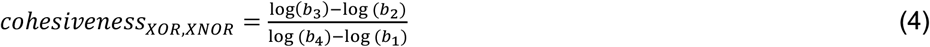

In each case, the equation was derived such that a perfectly functioning gate would approach a cohesiveness value of 1. We plotted each gate with cohesiveness versus dynamic range (Figure 3f). We predicted that visualizing the data in this manner would help us to visualize value and variance of ß and help to determine whether ß could be used as a single-descriptor of gate performance. Interestingly, in several cases we found papers where authors self-categorized specific gates as “good” or “bad” and we noted these annotations along with the regulation mechanisms (Figure S1, 3i).

### Fixed effect estimates are descriptors of logic gate performance

To analyze the application space and conceptualize representative gates, we performed unsupervised machine learning to cluster the data. Specifically, we aimed to break up the data such that we could visualize the “goodness” of existing published Boolean logic gates. We performed k-means clustering, as this algorithm is relatively simple but still widely applicable in many fields, including biology^59^, to parse these data. With our dataset, local maxima of the average silhouette scores were found for k = 3 and k = 6 clusters (Figure S1) and 6 clusters were selected for a more detailed separation of the data (Figure S1, S2, 3f).

We then simulated representative data for each of the clusters, following a Gaussian distribution using the centroids of each of the six clusters as mean responses for OR gate data (Figure 3b-e, g,h). A variance of 10% of the mean was chosen based on responses gathered from our nested repressor gate evaluation (discussed later). Each of these simulated responses (Figure 3b-e, g,h) are representations of data observed during our review of previous literature. Finally, we fit the data with the proposed linear mixed effects model (Equation 1) to obtain an estimate for the fixed effect ß and used a general linear hypothesis test to evaluate each gate for statistical significance.

Returning to the self-categorizations included in the data scrape, we noted that in the case of the clustering, the green (e.) cluster contained no author self-categorized “bad” data, and only self-categorized “good” data. This indicates that both our clustering and researcher intuition align for circuits with optimal performance. We then sought to examine whether ß could be used as a metric for gate performance. Interestingly, we only measured fixed effects statistically greater than zero (with a two-tailed hypothesis test) for two centroids: green (Figure 3e.) and indigo (Figure 3g.). To confirm this was not a statistical anomaly due to an unusual seed, we verified these results using Monte Carlo simulations (Figure S5-7). In the case of the indigo cluster (g.), which was also statistically significant, only one self-categorized “bad” point was present, and the primary categorization was “good”. Using specifications for commercially available CMOS electrical logic gates, we also simulated outputs for each logic (OR, NOR, AND, and NAND), and treated to the same data processing as the published genetic gates. These outputs were also placed in the green (e.) cluster, further validating the machine learning clustering, and ß as a metric of success. Importantly, when we examined the data for purple (c.) and yellow (d.) clusters, we noted a mixed categorization (“good” and “bad”) as well as a ß-estimates that were not significantly different than zero. We imagine that this high dynamic range region with a lower relative cohesiveness score is responsible for this mixed categorization yielding a high variance of the ß-estimate. Some cases may indicate a misclassification of logic, or a different biological phenomenon than originally designed. Broadly, we determined that the parameter ß is a representative of gate behavior where a higher estimate with lower variance is associated with improved gate performance (Figure 3j). In cases with a large of variance, the ß estimate is associated with a lower cohesiveness. Our analysis shows that researchers are adept at self-categorizing gates with very low or very high cohesiveness and ß values; but that intuition is less reliable for intermediate values (Figure 3c-3d.). Together, this analysis highlights the need for more rigorous statistical characterization of synthetic genetic circuits.

### Monte Carlo evaluation of mixed effects model for logic gate performance and sample size

To further assess the estimation of the fixed effects parameter, ß, as a holistic descriptor of gate performance, we evaluated whether the mixed effects model would return an unbiased estimate of the true fixed effect ß. Published biological datasets often contain low replicates per input group and have a low number of input groups (random effects factor levels), which can affect the linear mixed model fit and parameter estimation. We aimed to determine the model’s ability to estimate ß when provided with different sample sizes. For each gate, we simulated Gaussian data with a true fixed effect difference (ON-OFF) of 0.45 (a.u.) and a sample size of 3, 5, or 30 for gates OR, NOR, AND, and NAND (Figure 4), and XOR, XNOR, IMPLY, and NIMPLY (Figure S3). We chose these three sample sizes because both 3 and 5 are common samples sizes for biologists (“biological triplicate”) and 30 because it is a well-known sample size threshold in statistics. Then, we used Monte Carlo simulations for a linear mixed effects model to determine if the estimation exhibited bias or if it would converge on the true fixed effect. In each sample size, the results converged on 0.45 (a.u.) (Figure 4b, d, f, h, Figure S8).

**Figure 4.**
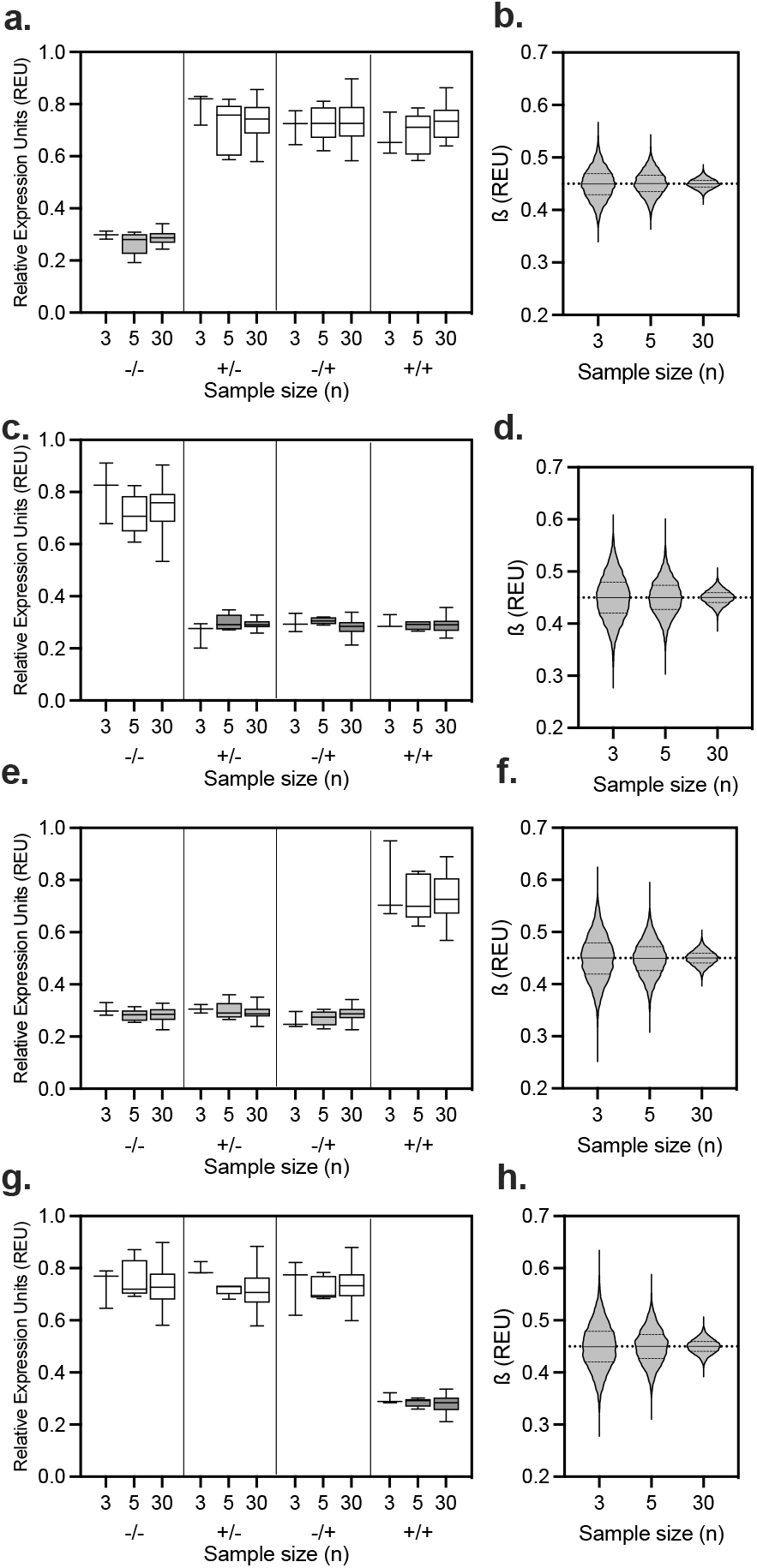
Model validation with given ß compared to estimate using mixed effects model. **a**. Example of simulated data for an “OR” gate output simulated from a Gaussian distribution with ß = 0.45 for a sample size of 3, 5 and 30 with a variance estimated as 10% of the mean. **b**. Monte Carlo simulations showing point estimates of ß from a model fit using data simulated from the same distribution as in **a**., ran 10,000 times for each sample size. The dotted line behind the data represents the true parameter value ß = 0.45, while the solid lines (from top to bottom) represent the top quartile, mean, and bottom quartile of the ß estimates. As sample size increases, point estimates of ß converge to the true value. **c. e**. and **g**. represent the same process applied to the NOR, AND, and NAND gates respectively with **d. f**. and **h**. showing their respective Monte Carlo simulations.

We occasionally observed a singular model fit^60^, where some of the variance-covariance matrix was estimated to be exactly zero^55^. This happened for both real and simulated data when fitting the model to data with few observations, small sample sizes, and few random effects; however, it does not affect the power of the statistical test (Figure S5-7) nor bias the point estimates of ß (Figure S8). Additionally, the linear mixed-effects model uses a homogeneity of variance, which our simulated data violates by assuming a variance of 10% of the mean. Nevertheless, we found that this violation does not bias the model nor reduce its power. Our Monte Carlo simulations prove within a multi-input group with a shared output (*i*.*e*., the “ON” state in an “OR” gate) that variations within these input groups can be attributed to the random effect. For well-performing gates, (Figure 3e, 3g, S5) with low to moderate sample variance (10-30% of mean), a sample size of n = 3 provides sufficient power. For mixed-performance gates with low sample variance (10% of mean, Figure 3h, S6) or well performing gates with higher sample variance (Figure 3e, 3g, 7, variance >50% of mean), increasing sample size will significantly increase the power of the statistical test. For poorly performing gates (Figure 3b, 3c, 3d, S7), regardless of population variance, an increase in sample size will not affect power.

### The fixed effects estimate ß correlates to biological gate performance in nested repressor OR gate

Finally, we evaluated ß as a descriptor of real biological data and examined its potential use in the forward design of genetic circuits. We chose a set of nested repressor OR gates controlling expression of a fluorescent protein^32,34^ and modulated the relative gate performance by controlling the RBS strength of the TetR and LuxR repressors (Figure 5). The RBS site variants were created using the Salis Lab RBS calculator “*De novo* DNA” for predicted translation initiation rates and design^46^. For our base strain of pOR-*sfgfp* in *E. coli* DH5α, the strength of the RBS upstream of TetR was predicted to have a proportional translation initiation rate of 4,362 a.u., and the RBS regulating LuxR was predicted to have a proportional translation initiation rate of 9,519 a.u. (Figure 5). When the relative strength of the LuxR_RBS_ was decreased over 10-fold, a comparable logic function was achieved. However, a 2-fold reduction in the TetR_RBS_ resulted in a drastic change in function (Figure S4). Contrary to our original expectation, we measured poorer performance in the nested repressor when modulating only the TetR_RBS_. This result reinforces that characterization in a biological system is challenging to break into constitutive components in the same manner that electrical systems can be parsed. We hypothesized that this breakdown is due to both regulator proteins being under control of the same P_*LacIQ*_ constitutive promoter with no insulating DNA between regulation elements and suggests that the upstream RBS of the TetR protein is affecting the translation initiation of both proteins.

**Figure 5.**
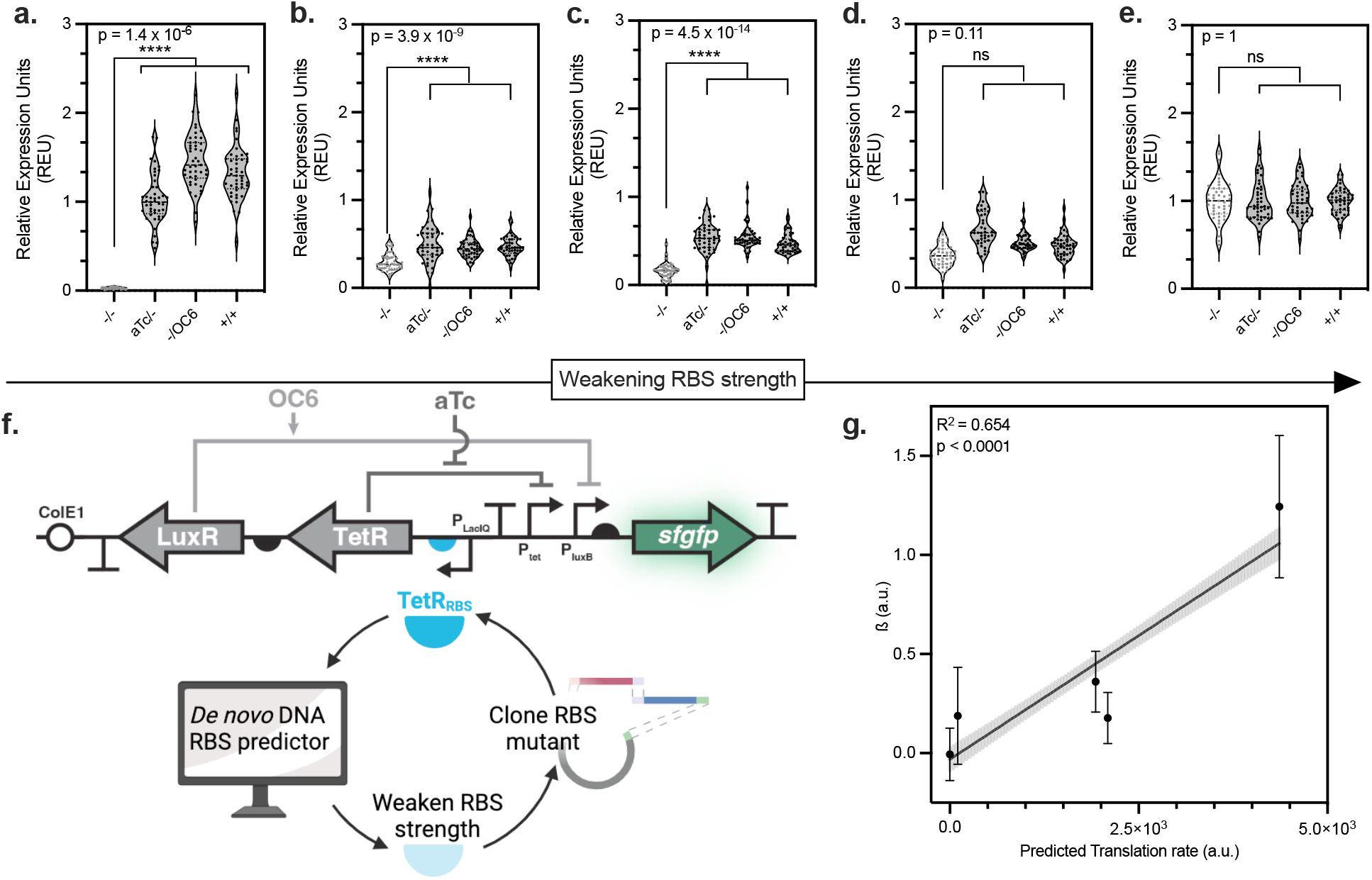
ß is a holistic indicator of OR gate function. **a**. Behavior of OR base gate in *E. coli* DH5α controlling expression of *sfgfp* and normalized by a constitutive strain lacking all promoter proteins. Data represents n=44. **b**. Behavior of OR gate weaker RBS (2,091 proportional relative translation rate) regulating LuxR in *E. coli* DH5α controlling expression of *sfgfp* and normalized by a constitutive strain lacking all promoter proteins (**e**.). Data represents n=44. **c**. Behavior of OR gate weaker RBS (1,931 proportional relative translation rate) regulating LuxR in *E. coli* DH5α controlling expression of *sfgfp* and normalized by a constitutive strain lacking all promoter proteins (**e**.). Data represents n=44. **d**. Behavior of OR gate weaker RBS (104 proportional relative translation rate) regulating LuxR in *E. coli* DH5α controlling expression of *sfgfp* and normalized by a constitutive strain lacking all promoter proteins (**e**). Data represents n=44. **e**. Behavior of OR gate constitutive strain lacking all promoter proteins. Data represents n=44. **f**. Schematic describing workflow iteration of the TetR_RBS_ design, test, build cycle. **g**. Correlation between the predicted translation rate of TetR and the ß estimate describing the fixed difference between ON and OFF gate function. Error bars represent the standard error of ß and the linear regression equation is y = 2.5e10^−4^ × −0.031.

Based on these results, we predicted that the RBS strength of TetR alone may control gate behavior and that this could be used as an indirect measure of performance. We utilized the design mode in *De novo* to create a low-strength TetR RBS (104 relative translation initiation rate) and two mid-strength TetR RBS (1,931 and 2,091 relative translation initiation rate) mutants. Finally, we synthesized a circuit lacking the TetR and LuxR regulator entirely, to create a constitutively expressed positive control by which we could benchmark gate performance. When quantifying the behavior of the OR-mutant strains, we normalized each output by the output of the constitutively expressed output to yield relative expression units (REU). We predicted that the stronger the RBS, the better performing the two-input Boolean gate and therefore higher estimate of ß. Qualitatively, we measured a drop-off in gate performance for the gates with weaker TetR RBS performance compared to the original RBS design (below a relative translation initiation rate of 4,000). Using a linear mixed effects model to examine each of the five OR gates, we determined that the three highest RBS gates (Figure 5c-e) exhibited statistically significant gate performance using a generalized linear hypothesis test (H_0_: ß=0). Leveraging the predicted relative translation rate of the TetR repressor as an indirect measure of gate performance we observed good agreement between gate performance and ß with a statistically significant slope (p<0.0001, R^2^=0.654) for a linear regression (Figure 5b). Together, these data suggest that ß can serve as an indicator of gate performance to direct design and assess gate success where a higher ß correlates to a better performing gate (Figure 5b).

## Discussion

We propose a linear mixed effects model for analyzing multi-input synthetic gene circuits. The fixed effect parameter, ß, serves as a holistic descriptor of gate performance by accounting for both dynamic range and agreement within states. We first evaluated the linear mixed effects model and the performance of ß specifically on a set of previously published synthetic gene circuits. From our evaluation of these circuits, we determined that there is no inherent correlation between the mechanism of genetic regulation, type of logic (AND, NAND, etc.), and gate performance (Figure S2). Highlighting the need for a quantitative descriptor under these conditions, the ß estimate obtained from the model was used to statistically analyze biological Boolean gate performance. Estimates of ß statistically different from zero (equivalent to rejection of H_0_: ß=0) indicate that there are two distinct states being accessed. As expected, we also found that researchers have an intuitive understanding of ‘good’ versus ‘poor’ performance for a given synthetic gene circuit. However, this intuition starts to break down in cases where one ON (or OFF) condition is substantially different from others belonging to the same state (e.g., Figure 3d). While we applied the linear mixed effects model solely for 2-input/1-output Boolean logic, in principle, it could be expanded to account for multi-input^47^ and multi-output states. Finally, in all cases, we utilized frequentist statistics instead of Bayesian statistics because many genetic logic gate sample sizes are small for Bayesian statistical analyses, which require many more design choices in their implementation^61^. However, we note that similar analyses could be completed utilizing Bayesian statistics.

A significant challenge in assessing the success of synthetic gene circuits is the number of potential output signal types (or states) and their dependence on experimental conditions. Nevertheless, we envision that maximizing ß may prove useful in determining what analytical technique is most appropriate for characterizing a given synthetic gene circuit. Often application and evaluation of synthetic gene circuits occur in a population of cells (*i*.*e*., a liquid culture) with batch population measurements (*i*.*e*., plate reader). If a researcher is interested in assessing the performance of new genetic regulation technology, then flow cytometry and other single-cell measurements may be better suited to evaluate success compared to batch measurements of a population. Analyzing expression on a single-cell basis allows researchers to capture whether signal production is heterogeneous within the population (representing a bimodal or other non-unimodal response), or if the response is homogenous through the population. Establishing whether the response is hetero- or homogenous may be key in understanding potential applications as well as evaluating gate efficacy. However, batch measurements of cellular populations are sufficient for certain applications such as biosensors or control over material properties (color, stiffness, etc.), as well as in situations where a phenotype is difficult to determine at the single-cell level. Regardless of the method of measurement, a linear mixed effects model can be utilized to analyze the outcome.

For the most in-depth analysis of synthetic gene circuits and to compare across modes of regulation, we recommend (1) normalizing data to a constitutively “ON” state (REU), (2) continuing to publish self-described “bad” gates, and (3) reporting individual replicates and raw data. To facilitate the analysis of these circuits, raw or normalized data can be analyzed using the ‘lme4’ package^56^ in R for a frequentist mixed effects model. The output will provide estimations for ß which can then be tested for statistical significance. Example code, as well as tutorials and templates for how to calculate ß under a variety of different conditions, can be found in the supplementary information (Table S6 - S10). We have also provided an online calculator that can be used for calculating ß, cohesiveness, and dynamic range of 2-input gates. This online calculator lets the user select the gate type (currently limited to 2-input gates) and input raw data. It will then complete the analysis based on user input (https://boolstats.fly.dev/). When examining ideal gate behavior, we suggest that the output for each output state should strive for input invariance, meaning all ON states should yield the same degree of “ON”, in order to be categorized as an effective Boolean logic gate. Intermediate, non-binary states—while incredibly useful—are not binary Boolean outputs but rather represent a different class of logic where digital combinations of input inducers achieve analog expression outputs^16^. If intermediate states are present, this will be reflected in either a low ß value or a ß value with a high amount of variance. These analog outputs can be applied for more advanced computational tasks including memory, which is a feature of biological logic gates that is still underutilized^16^.

Interestingly, while biologists have taken inspiration from electrical and computer engineering in the creation of biological Boolean logic gates, there is an inverse phenomenon occurring with advances in non-traditional forms of computing that emulate living systems. For example, neuromorphic devices are high-performing and low power computation systems that are being investigated as an alternative to Von Neumann architectures^62^. Modern computers, as examples for Von Neumann architecture-based devices, have their voltage (or current) levels for binary logic shaped jointly by the commercial and United States military sectors, with key considerations given to switching speed, power consumption, and device reliability^63^. However, neuromorphic computing devices are now encountering challenges that biologists often come across including how to determine what is appropriately binary when measuring an analog output and how to discretize the inherently analog behavior of neurons for representation in the digital world^64,65^. To address some of these challenges in synthetic biology, we propose the metric ß to provide a quantitative description of genetic circuit success. In addition to functioning as a tool for statistical analysis, ß can also assist with building higher order logic, troubleshooting, or applying genetic circuits in applications that require rigorous statistical evaluation (*i*.*e*., healthcare). Better estimates of ß (higher with a lower variance, and statistical difference from zero) are characteristic of improved gate function. This holistic measurement also allows for performance comparisons across gate types (OR vs AND, etc.) which may yield valuable insight about underlying biological regulatory mechanisms or potential applications. In total, ß may be used to discern subtle differences in genetic circuit behavior, which may allude researcher intuition, and reveal valuable mechanistic insights into Boolean gate performance to help drive the next generation of applications for synthetic gene circuits.

## Materials and Methods

### Materials

Kanamycin sulfate (C18H38N4O15S, Growcells), anhydrotetracycline hydrochloride (aTc, Sigma-Aldrich), 3-oxohexanoyl-homoserine lactone (OC6, Sigma-Aldrich), NEB Gibson Assembly Master Mix (NEB E2611), NEB 5-alpha *E. coli* (NEB C2987). All media components were autoclaved or sterilized using 0.2 μm PES filters.

### Model description

The general form of a linear mixed-effects model using Bates’ notation^56^ takes the following form:

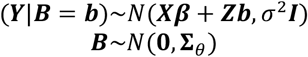

In the most general case, ***B*** is a q-dimensional random effects vector with a multivariate normal distribution around 0 and variance-covariance matrix **Σ**_*θ*_, and ***Y*** is an n-dimensional vector of the response data. The response observations ***Y*** at a fixed ***B*** (***B*** = ***b***) are all assumed to be independent with a constant variance *σ*^2^; ***I*** describes the *n* _×_ *n* identity matrix. The conditional distribution of ***Y*** given ***B*** = ***b*** is multivariate normal and has an expected value of ***Xβ*** + ***Zb. β*** is a p-dimensional fixed effects vector which is conceptually identical to ***β*** in a simple linear regression model, while ***X*** and ***Z*** correspond to the design matrices with dimensions *n* _×_ *p* and *n* _×_ *q*, respectively.

For the specific case of modeling of a two-input Boolean logic gate with a linear-mixed effects model, the model equation can be decomposed to the following:

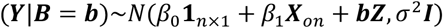

In this formulation, ***X***_*on*_ is a *n* _×_ 1 indicator variable with each component taking a value of 0 or 1, corresponding to an OFF or ON state, respectively, based on expected gate behavior. **1**_*n*×1_ is a vector of all ones, with dimensions *n* _×_ 1. The design matrix ***Z*** has dimensions *n* _×_ 4, corresponding to the four input states for two-input Boolean logic gates (*q* = 4). In the design matrix ***Z***, the component at position *i, q* takes a value of 1 if observation *i* is a member of input group *q* and 0 otherwise. This model is formulated for the lme4 package in R with the following syntax: “Results ~ on + (1|group)” where ‘on’ refers to the fixed effect *β*_1_ and ‘group’ represents the random effect (input grouping).

### Biological Boolean Logic Gate space and data scrape

22 papers^7,8,14,16,21–23,25,27,34,35,42,43,47,66–73^ were analyzed, containing 147 graphs in total. Of these graphs, data were scraped using automeris.io to determine the average of the output for a specific input group. Papers were chosen to represent a wide array of technologies available for biological Boolean logic. The means were processed by averaging the “ON” values for each gate and normalizing each gate output by the average of the ON values. Then, calculations were performed on the normalized data to determine the average of the ON outputs (which, by definition, is equal to 1) minus the average of the OFF outputs. The cohesiveness score for each gate was determined using equations 2-4. Prior to machine learning, 3 outliers were removed, yielding 144 analyzed gates.

The k-means algorithm was implemented with the Scikit-learn module in Python^74^ after standard centering and scaling. To select the number of clusters for k-means, we iteratively computed the average silhouette score from k = 2 to k = 10 clusters. We observed a local maxima of average silhouette score for k = 3 and k = 6 clusters (Figure S1) and selected k = 6 for increased precision. Code is available in Supplementary Table S7, S8. Voronoi lines separating clusters were identified via the SciPy module in Python^75^.

### Simulations of gate responses

Using the centroid points determined from k-means clustering, a system of equations was solved using Python to determine the overall gate response for an “OR” gate that satisfy the parameters. Code is available in Supplementary information S8. Then, data simulation was performed in R for these average responses of the centroid as determined by the systems of equations with a variance of 10% of the mean and data was simulated assuming a normal distribution. For every simulation, the random seed was set to ‘2024’ for code reproducibility. The code is described in Table S8. For the Monte-Carlo estimation of ß in R, a for loop was used to simulate and then estimate ß ten thousand times for each case. The code is described in Table S11.

### Gibson Cloning of TetR RBS library

Constructs were designed such that a single PCR reaction with proper Gibson Cloning primers would yield overhangs corresponding to the RBS site of interest. Each primer contained identical overlap regions on the starting plasmid. PCR Conditions for Q5 High-fidelity PCR reaction (from NEB) are found in Supplementary Table S4, S5. After PCR amplification, 8.3 μL was removed from each reaction (for gel confirmation), and 1 μL of DPNI was added for a 1 h incubation at 37°C. A PCR clean-up was performed using a Quiagen PCR purification kit, and a Gibson reaction with 0.045 pmols of the PCR product with 2X High-Fidelity Gibson Master Mix (NEB) was incubated at 50°C for one hour. DNA was introduced using a heat shock protocol and plated onto 50 μg/mL kanamycin containing LB agar plates. Sequence confirmation from Plasmidsaurus was used to confirm plasmid identity and introduction of new RBS site. All plasmids are described in Supplementary Table S2 with links to full plasmid sequencing in Supplementary Table S12. Primers are outlined in Appendix S3.

### Bacteria Strains and Culture

Bacterial strains and plasmids are listed in Table S7.1. Cultures were prepared from bacterial stocks stored in 20% glycerol at −80 °C streaked onto LB agar plates (for wild-type and knockout strains) and grown overnight at 37 °C for *E. coli*. Overnight cultures were grown by picking single colonies and inoculating LB broth. Plasmid harboring strains were grown with the addition of 50 μg/mL of kanamycin diluted from a 1000× stock in water. OD_600_ was measured using a NanoDrop 2000C spectrophotometer.

### Gate evaluation and fluorescence

OR gates were evaluated by inoculating into a 96-well plate containing 200 μL of LB with 50 μg/mL kanamycin per well. Each plate contained a constitutive *sfgfp* strain in *E. coli* K12 which was used as a standard to determine plate-to-plate variability. After growing for 16 h, 10 μL from each well was diluted into a new plate containing LB, kanamycin, and the inducer of interest. The same overnight cultures were used to inoculate into each induction condition making a pairwise comparison. These reactions were sealed and allowed to grow for 16-18h. Then fluorescence and OD_600_ were collected from each sample. REU was determined by first background subtracting, then normalizing the sample by the constitutive control in *E. coli* DH5α OR gate lacking all repressor proteins.

## Data Availability

Data supporting the findings of this study are available within the Supplementary File and through the Texas Data Repository (https://doi.org/10.18738/T8/CIESXN). Biological materials are available upon request to B.K.Keitz.

## Acknowledgments and Funding Sources

This research was financially supported by the Welch Foundation (grant F-1929, B.K.K.), the National Institutes of Health under award number R35GM133640 (B.K.K.), a National Science Foundation (NSF) CAREER award (grant 1944334, B.K.K.), the Air Force Office of Scientific Research under award number FA9550-20-1-0088 (B.K.K.) and the National Science Foundation (CBET-2133661) (HSA). GP and SMC were funded by were supported through NSF Graduate Research Fellowships (program award DGE-1610403). C.O.W. was supported by the Jane and Roland Blumberg Centennial Professorship in Molecular Evolution. We would like to thank Alexis J. Holwerda for her cloning expertise. Schematics were created using BioRender.com, graphs were created in Prism GraphPad and Boolean logic statistics were run in R.

