## Supplemental Information for "A Linear Mixed Effects Model for Evaluating Synthetic Gene Circuits"

**This PDF file includes:**

Supporting text
Figures S1 to S9
Tables S1 to S12

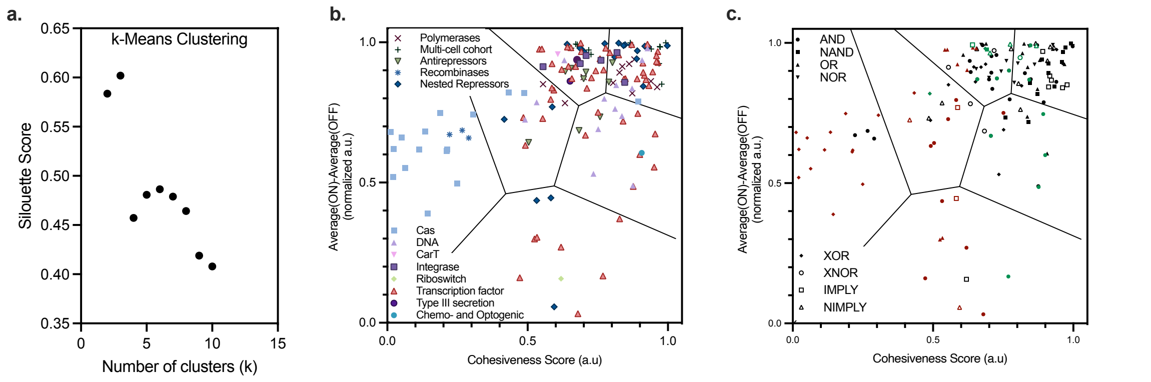


**Figure S1. a**. Silhouette score for K-means clustering. Scrape Analysis **b.** data reprinted from Figure 3f. overlaid with broad-categorization of biological regulation machinery. **c.** Data reprinted from Figure 3f. overlaid both with gate type (Boolean Logic) and which points are self-categorized as "good" (green) or "bad" (maroon).

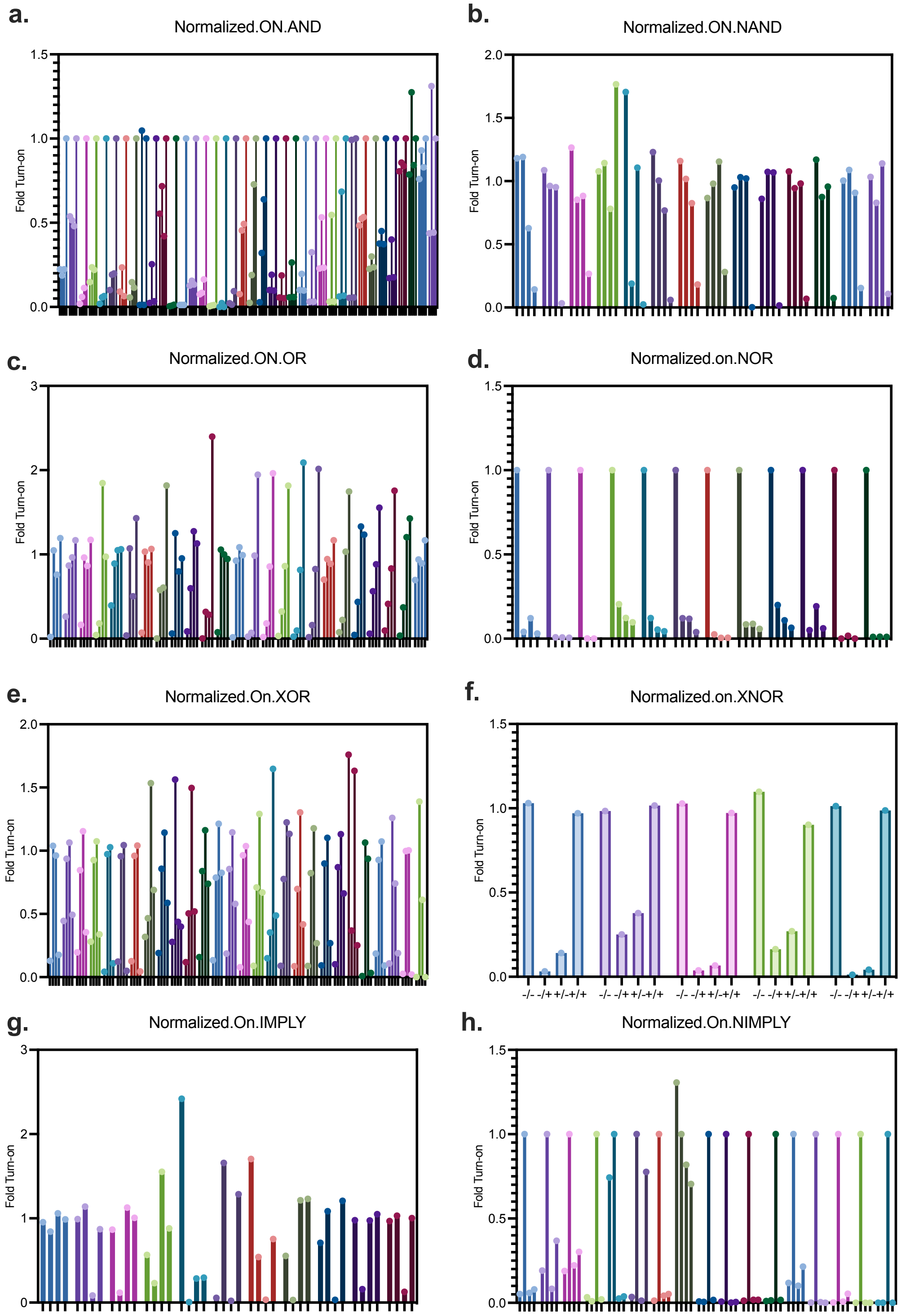


**Figure S2. Raw input data for each figure 3f.** Each color cluster represents the mean responses of a single logic gate. (-/-, -/+, +/-, +/+) for Boolean logic gates: **a.** AND **b.** NAND **c.** OR **d.** NOR, **e.** XOR, **f.** XNOR, **g.** IMPLY, **h.** NIMPLY.

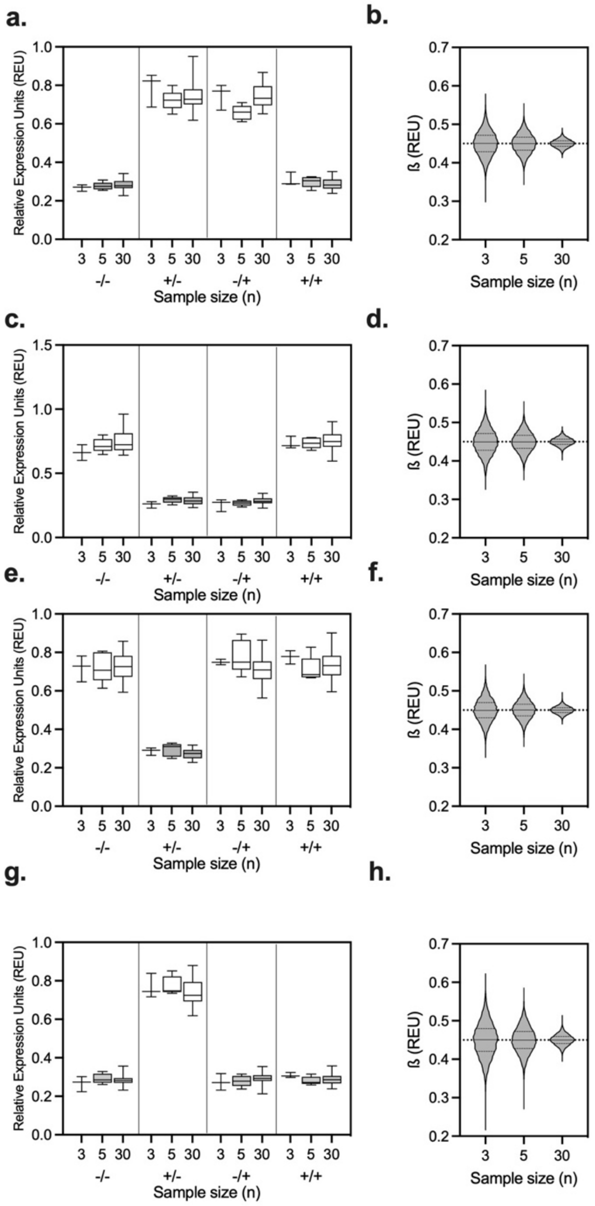


**Figure S3. Model validation with given ß compared to estimate using mixed effects model. a.** Simulated data for an "XOR" gate output simulated with ß = 0.45 for a sample size of 3, 5 and 30 given a 10% variance in each mean. **b.** Monte Carlo simulations for ß estimates for **a.** run 10,000 times for each sample size. Dotted line represents ß = 0.45 which was used to simulate the data in **a.** and shows good convergence in Monte Carlo simulation with tighter estimations for a higher sample size. **c. e.** and **g.** Represent the same process applied to the XNOR, NIMPLY and IMPLY gates respectively with **d. f.** and **h.** showing their respective Monte Carlo simulations.

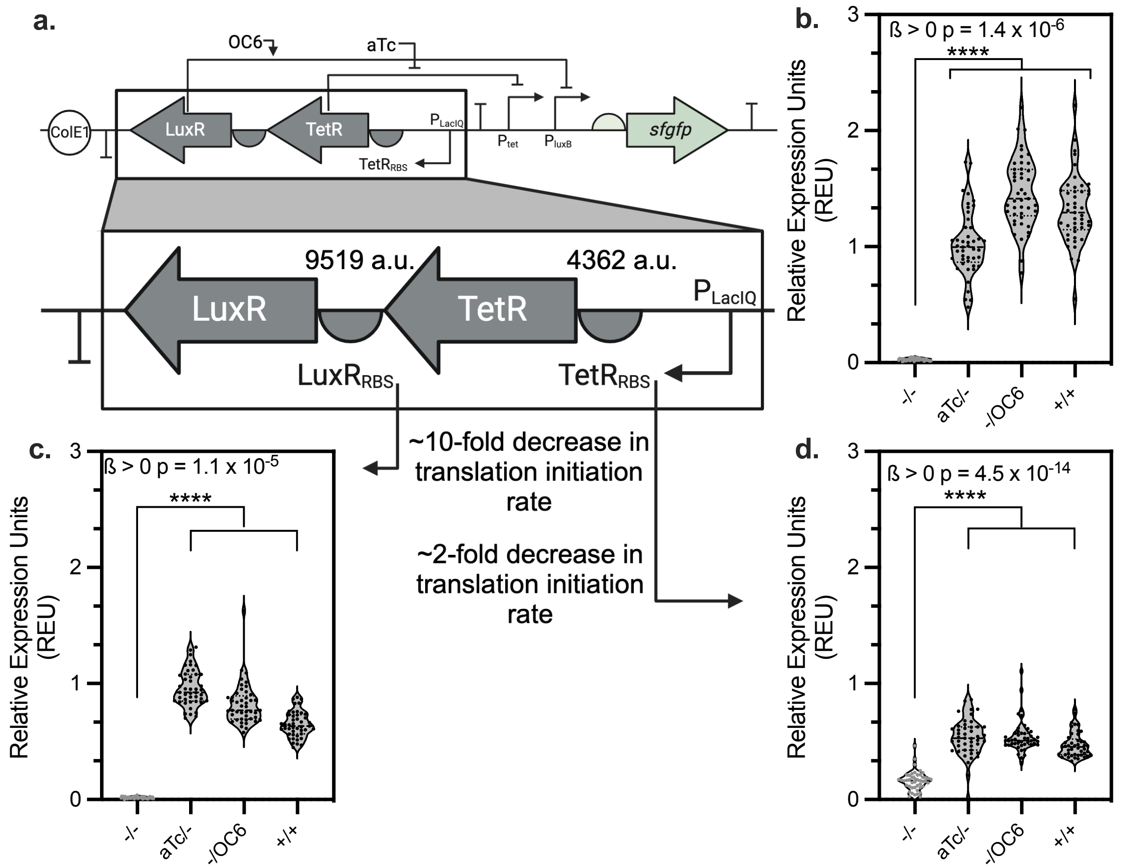


**Figure S4. Nested repressor regulation of $TetR_{RBS}$ controls expression of both LuxR and TetR regulatory proteins due to lack of insulating DNA between sequences. a**. Nested repressor OR logic gate with call out highlighting the regulatory binding proteins TetR and LuxR and their RBS. **b**. Behavior of OR base gate in *E. coli* DH5ɑ controlling expression of *sfgfp* and normalized by a constitutive strain lacking all promoter proteins. Data represents *n*=44. Data in b. is reprinted from Figure 5 for clarity. **c.** Behavior of OR gate with a 10X weaker RBS regulating LuxR in *E. coli* DH5ɑ controlling expression of *sfgfp* and normalized by a constitutive strain lacking all promoter proteins. Data represents *n*=44. **d.** Behavior of OR gate with a 2X weaker RBS regulating TetR in *E. coli* DH5ɑ controlling expression of *sfgfp* and normalized by a constitutive strain lacking all promoter proteins. Data represents *n*=44. Data in d. is reprinted from Figure 5 for clarity.

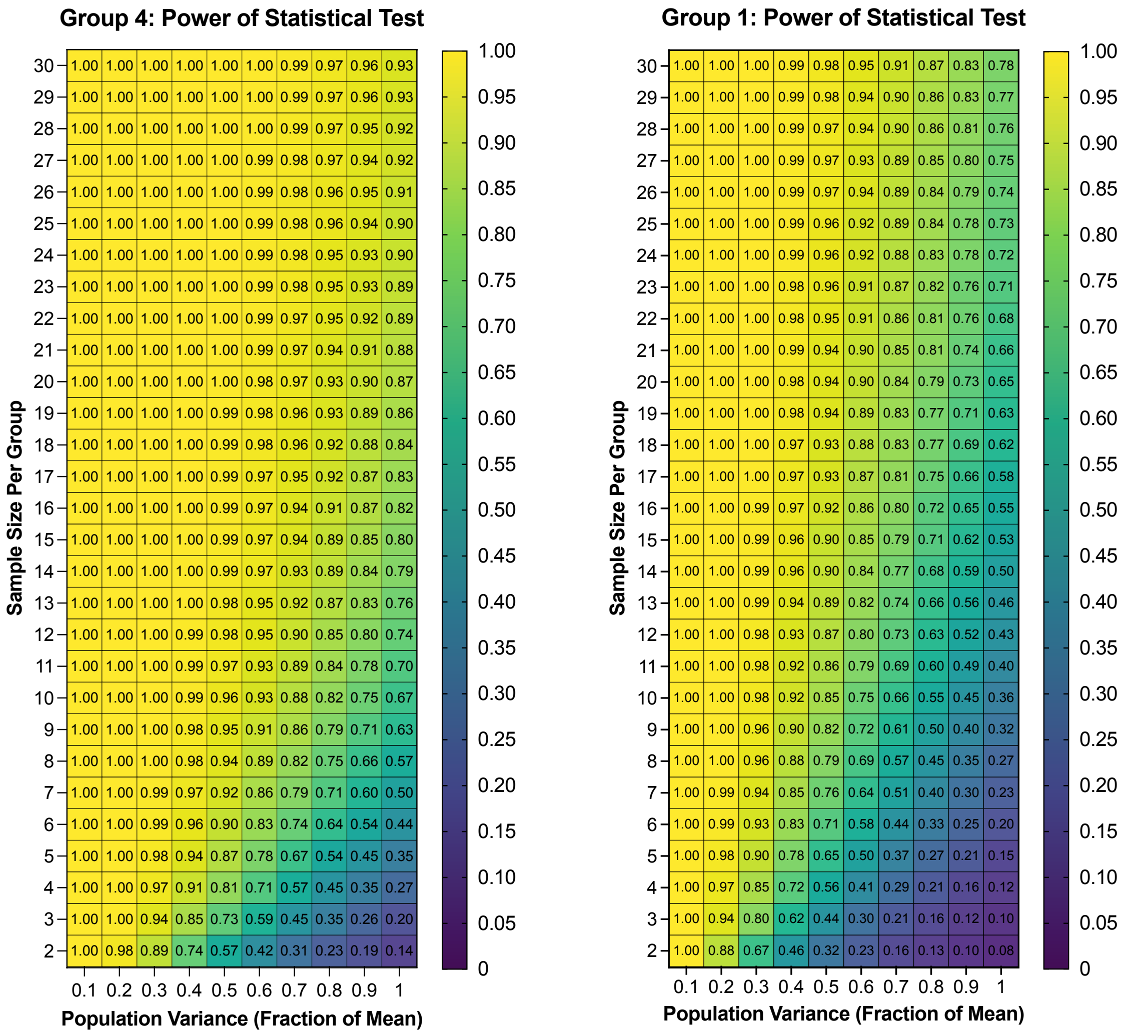


**Figure S5.** Power analysis of the well-performing gates. Group 4 is from Figure 3e. and Group 1 is from Figure 3g.

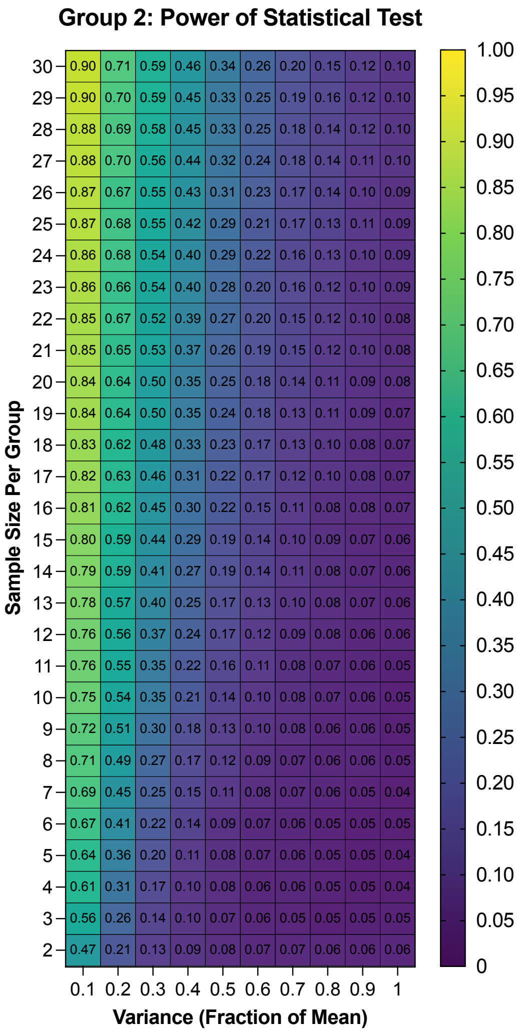


**Figure S6.** Power analysis of the middle-performing gate, with low dynamic range. Group 2 is from Figure 3h.

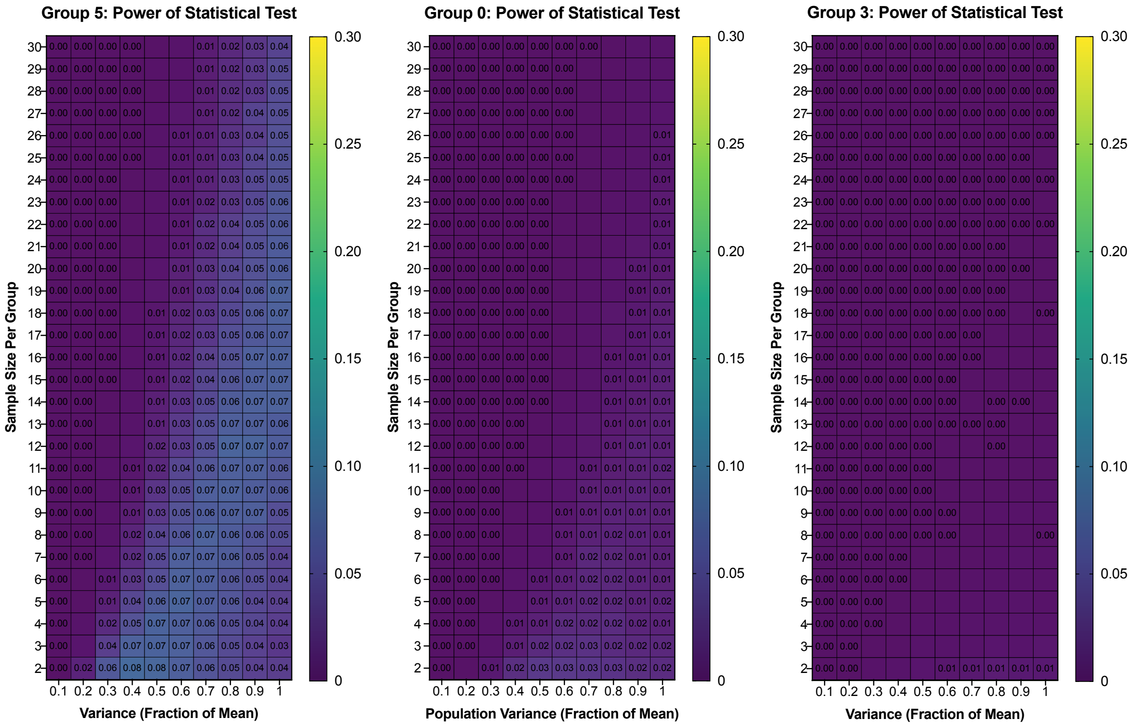


**Figure S7.** Power analysis of the poorly-performing gates. Group 5 is from Figure 3d, Group 0 is from Figure 3c. and Group 3 is from Figure 3b.

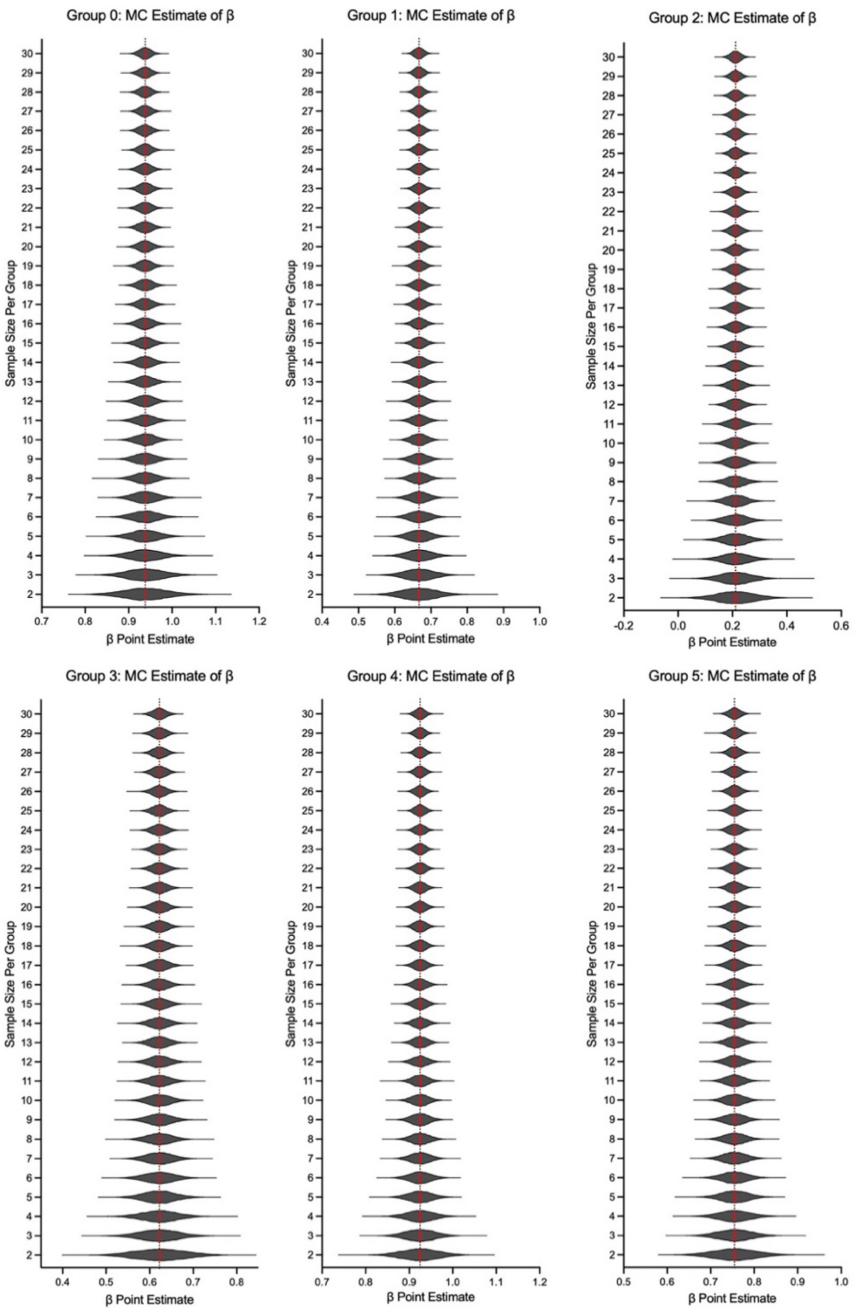


**Figure S8.** β point estimates are unbiased estimators of the true average(ON) – average(OFF) performance for the six simulated k-means gates. At a simulated variance corresponding to 10% of the true mean for each group, and from 2 to 30 samples per group, the mean of β point estimates (solid red line) is equal to the true average(ON) – average(OFF) (dashed black line). Group 0 is from Figure 3c., Group 1 is from Figure 3g., Group 2 is from Figure 3h., Group 3 is from Figure 3b., Group 4 is from Figure 3e,. and Group 5 is from Figure 3d.

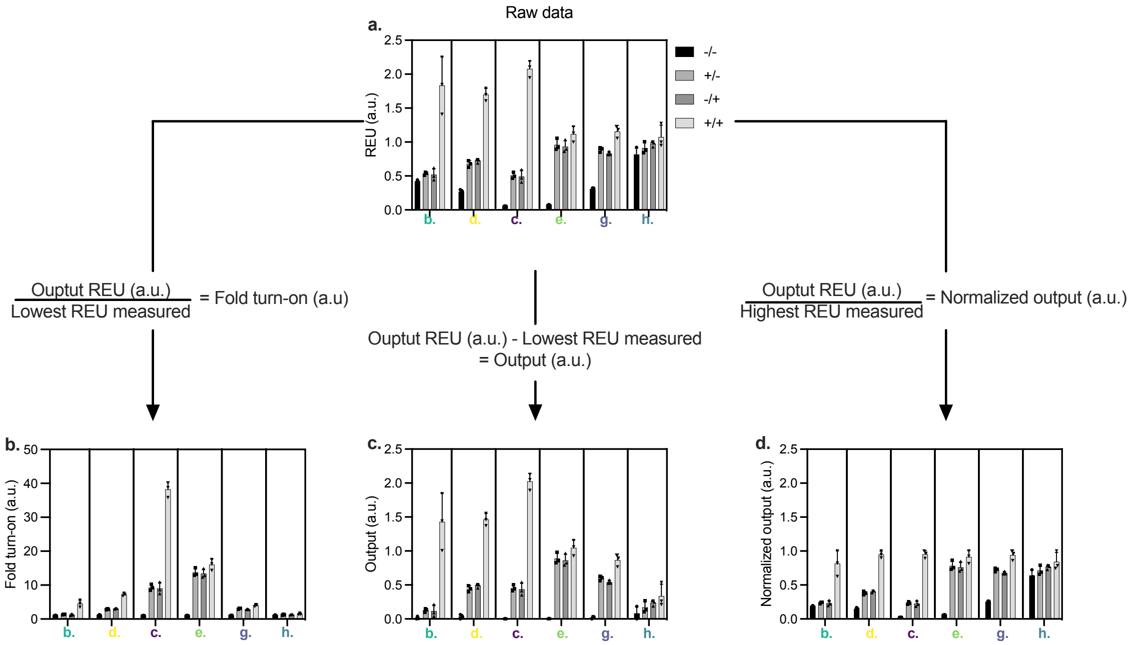


**Figure S9.** Simulated data from Figure 3 represented in different data pre-processing. **a.** Simulated "Raw" data from Figure 3. **b.** Data from (**a**.) normalized by dividing the lowest measured REU to get "Fold turn-on". **c.** Data from (**a**.) processed by background subtracting lowest measured REU. **d.** Data from (**a.**) normalized by dividing the highest measured REU.

**Supplementary Table 1.** Boolean Logic Gates

| Gate | -/- | -/+ | +/- | +/+ |
|---|---|---|---|---|
| OR | 0 | 1 | 1 | 1 |
| NOR | 1 | 0 | 0 | 0 |
| AND | 0 | 0 | 0 | 1 |
| NAND | 1 | 1 | 1 | 0 |
| XOR | 0 | 1 | 1 | 0 |
| XNOR | 1 | 0 | 0 | 1 |
| IMPLY | 0 | 1 | 0 | 0 |
| IMPLY | 0 | 0 | 1 | 0 |
| NIMPLY | 1 | 0 | 1 | 1 |
| NIMPLY | 1 | 1 | 0 | 1 |

**Supplementary Table 2**. Bacterial strains and plasmids used in this study.

| Strain or plasmid | Description/Genotype | Reference or source |
|---|---|---|
| *E. coli DH5α* | *LacZ*ΔM15, *rec*A1 mutation, *endA* mutation, high efficiency | Invitrogen C2987 |
| *E. coli DH5α* + pOR_sfgfp | + pOR_sfgfp | Graham, Partipilo et al. 2024 |
| *E. coli DH5α* + pOR_med | + pOR_med | This work |
| *E. coli DH5α* + pOR_med.low | + pOR_med.low | This work |
| *E. coli DH5α* + pOR_low | + pOR_low | This work |
| *E. coli DH5α* + pOR_Constitutive | + pOR_Constitutive | This work |
| *E. coli DH5α* + pOR_LuxR1k | + pOR_LuxR1k | This work |
| Plasmids | | |
| + pOR_sfgfp | TetR/LuxR nested repressors “OR” logic with TetR RBS strength of 4362.58 relative translation rate | Graham, Partipilo et al. 2024 |
| + pOR_med | TetR/LuxR nested repressors “OR” logic with TetR RBS strength of 2091.22 relative translation rate | This work |
| + pOR_med.low | TetR/LuxR nested repressors “OR” logic with TetR RBS strength of 1931.42 relative translation rate | This work |
| + pOR_low | TetR/LuxR nested repressors “OR” logic with TetR RBS strength of 104.58 relative translation rate | This work |
| + pOR_Constitutive | pOR logic lacking both TetR/LuxR nested repressors yielding constitutively expressed *sfgfp* | This work |
| + pOR_LuxR1k | TetR/LuxR nested repressors “OR” logic with LuxR RBS strength of 915.63 relative translation rate | This work |

**Supplementary Table 3.** Genetic parts/sequences used to construct the plasmids in this study.

| Genetic Part | DNA Sequence (5’ to 3’) |
|---|---|
| Ribosome Binding Sites | |
| TetR (pOR_*sfgfp*) | CTATGGACTATGTTTTCACACAGGAAAGGCCTCG |
| TetR (pOR_med) | GCGCATCTCGTATTCCGGCCATTAATTAT |
| TetR (pOR_med.low) | TCCAACAGAATACCGATAAGCTTCCCTTA |
| TetR (pOR_low) | TTATTTACATATTTACATACGATAAT |
| LuxR (pOR_LuxR1k) | TAAAACTTCACTTAATCTACACTCT |
| Gibson Cloning Primers | |
| TetR (pOR_med) FWD | GCGCATCTCGTATTCCGGCCATTAATTATGTCTCGTTTAGATAAATCTAAAG |
| TetR (pOR_med) REV | ATAATTAATGGCCGGAATACGAGATGCGCGGGCGCTATCATGCCATAC |
| TetR (pOR_med.low) FWD | TCCAACAGAATACCGATAAGCTTCCCTTAATGTCTCGTTTAGATAAATCTAAAG |
| TetR (pOR_med.low) REV | TAAGGGAAGCTTATCGGTATTCTGTTGGAGGGCGCTATCATGCCATAC |
| TetR (pOR_low) FWD | TTATTTACATATTTACATACGATAATATGTCTCGTTTAGATAAATCTAAAG |
| TetR (pOR_low) REV | ATTATCGTATGTAAATATGTAAATAAGGGCGCTATCATGCCATAC |
| Constitutive FWD | ACCTCGAGTCTCGGTACCAAATTCCAGAA |
| Constitutive REV | TTGGTACCGAGACTCGAGGTGAAGACGAAAGG |
| LuxR (pOR_LuxR1k) FWD | TAAAACTTCACTTAATCTACACTCTATGAAAAACATAAATGCCGAC |
| LuxR (pOR_LuxR1k) FWD | AGAGTGTAGATTAAGTGAAGTTTTATTAAGAACCAGATTCACATTTTAA |

**Supplementary Table 4.** PCR Components

| Component | 50 µl Reaction | Final Concentration |
|---|---|---|
| Q5 High-Fidelity 2X Master Mix | 25 µL | 1X |
| 10 µM Forward Primer | 2.5 µL | 0.5 µM |
| 10 µM Reverse Primer | 2.5 µL | 0.5 µM |
| Template DNA | 1.5 µL | < 1,000 ng |
| Nuclease-Free Water | 18.5 µL | |

**Supplementary Table 5.** PCR Conditions

| STEP | TEMP | TIME |
|---|---|---|
| Initial Denaturation | 98°C | 30 seconds |
| 20 cycles | 98°C | 10 seconds |
| | 54 + 0.5 °C/cycle | 20 seconds |
| | 72°C | 2 minutes |
| 10 cycles | 98°C | 10 seconds |
| | 64°C for 10 cycles | 20 seconds |
| | 72°C | 2 minutes |
| Final Extension | 72°C | 2 minutes |
| Hold | 4–10°C | |

**Table S6. R function to simulate Boolean gate data from a normal distribution.**

```
simulate_data <- function(working.params.df, sample.size, logic, fileNamePrefix, write.file = TRUE) {
  working.state.names <- row.names(working.params.df)
  working.df <- data.frame() # initaliize

  for (i in 1:length(working.state.names)){
    for (j in 1:length(sample.size)){

      state <- working.state.names[i]
      n <- sample.size[j]
      var_1 <- working.params.df[i,1] * 0.1
      var_2 <- working.params.df[i,2] * 0.1
      var_3 <- working.params.df[i,3] * 0.1
      var_4 <- working.params.df[i,4] * 0.1

      sim.1 <- rnorm(n, working.params.df[i,1],var_1)
      sim.2 <- rnorm(n, working.params.df[i,2],var_2)
      sim.3 <- rnorm(n, working.params.df[i,3],var_3)
      sim.4 <- rnorm(n, working.params.df[i,4],var_4)

      Results <- c(sim.1,sim.2,sim.3,sim.4)

      group <- as.factor(c(rep(1,n),rep(2,n),rep(3,n),rep(4,n)))
      on <- as.factor(c(rep(logic[1],n),rep(logic[2],n),rep(logic[3],n),rep(logic[4],n))
      cat(paste("Now doing state",state,"sample size",n,'.','\n'))
      working.df = data.frame(Results, on, group)

      if (write.file) {
        write.csv(working.df,paste0(fileNamePrefix,'_sample_size_',n,'_simulated_data.csv'))
      }
    }
  }
  return(working.df)
}
```

**Example use:**

```
setwd('/Users/)

set.seed(2024)
source("simulate_data.R")
library('lme4')
library('multcomp')

### SIMULATION ###
test.data.frame <- data.frame(t(c(0.1,0.8,0.8,1))) # data frame with mean
# responses to simulate, logic must match
row.names(test.data.frame) <- c('example_or_gate') # helpful for your own
# notation, or checking simulation progress

sample.size <- c(3,5) # number of samples per group
logic <- c(0,1,1,1) # expected

simulate_data(test.data.frame, sample.size, logic, "file_name_prefix") # writes
# two csvs with code in your working directory, and also returns dataframe
# with simulated data

### ANALYSIS ###
working.df <-
read.csv('file_name_prefix_sample_size_3_simulated_data.csv',header=TRUE,row.names=1)
working.lmer <- lmer(Results ~ on + (1|group), data = working.df)

print(confint(summary(glht(working.lmer,linfct=c('on==0')))))
  print(summary(glht(working.lmer,linfct=c('on==0'))))
```

**Table S7. R Script for Figure 3: k-means clustering, simulation, and statistics**

```
setwd('/Users/)

seed = 2024
set.seed(seed)
library('lme4')
library('effects')
library('broom')
library('simr')
library('multcomp')

source("simulate_data.R")

new.cond <- read.csv('20240806_data_k_6.csv',header=TRUE,row.names=1)
sample.size <- 3
or.logic <- c(0, 1, 1, 1)

for (i in 1:length(row.names(new.cond))){ # this loop simulates the data
  working.row.name = paste0('20240806_sim_data_k_6_',row.names(new.cond)[i])

  or.results.i <- simulate_data(new.cond[i,1:4], sample.size, or.logic, working.row.name)
}

for (i in 1:length(row.names(new.cond))){ # this loop does the statistics

  fileName =
paste0('20240806_sim_data_k_6_',row.names(new.cond)[i],'_sample_size_',sample.size,'_simulated_dat
a.csv')
  working.df <- read.csv(fileName, header=TRUE, row.names = 1)
  working.lmer <- lmer(Results ~ on + (1|group), data = working.df)

  cat(paste0('############ Reading filename ',fileName))
  print(confint(summary(glht(working.lmer,linfct=c('on==0')))))
  print(summary(glht(working.lmer,linfct=c('on==0'))))

}
```

**Table S8. Python Script for determining centroid values**

```
import numpy as np
from scipy.optimize import minimize

# Known value
x = #(1-b)

# Target values
target_log_ratio = #fromcentroid

# Objective function to minimize (difference from target log ratio)
def objective(vars):
w, y, z = vars
penalty = 1000 * np.abs(w + y + z - 3) # Penalty for violating the constraint
log_wy = np.log(w * y)
log_x2 = np.log(x**2)
log_z2 = np.log(z**2)
log_ratio = (log_wy - log_x2) / (log_z2 - log_x2)
return np.abs(log_ratio - target_log_ratio) + penalty

# Constraints
```

```
constraints = (
{'type': 'eq', 'fun': lambda vars: vars[0] + vars[1] + vars[2] - 3}, # w + y + z = 3
)

# Bounds for w, y, and z
bounds = [(0.1, 2), (0.1, 2), (0.1, 2)] # Adjust bounds if needed to meet x < w < y < z

# Initial guess
initial_guess = [1.0, 1.0, 1.0]

# Solve the optimization problem
result = minimize(objective, initial_guess, bounds=bounds, constraints=constraints)

# Extract results
w, y, z = result.x

w, y, z
```

**Table S9. Script for Figure 4 simulation, ß estimation, and statistics**

```
  setwd('/Users/)
  source("simulate_data.R")

  set.seed(2024)
  library('lme4')
  library('effects')
  library('broom')
  library('multcomp')

  sample.size <- c(3,5,30)

  new.cond <- read.csv('New.conditions.4.Sarah.csv',row.names=1,header=TRUE)

  and.logic <- c(0, 0, 0, 1)
  or.logic <- c(0, 1, 1, 1)
  nand.logic <- c(1, 1, 1, 0)
  nor.logic <- c(1, 0, 0, 0)
  xnor.logic <- c(1, 0, 0, 1)
  xor.logic <- c(0, 1, 1, 0)
  imply.logic <- c(1,0,1,1)
  nimply.logic <- c(0,1,0,0)
```

```
##### SIMULATES THE DATA #####

for (j in 1:length(sample.size)){
 size <- sample.size[j]
 i = 1
 or.results <- simulate_data(new.cond[i,1:4], size, or.logic,
paste0(row.names(new.cond)[i],'_sample_size_',size,'_simulated_data.csv'))
 i = 2
 nor.results <- simulate_data(new.cond[i,1:4], size, nor.logic,
paste0(row.names(new.cond)[i],'_sample_size_',size,'_simulated_data.csv'))
 i = 3
 and.results <- simulate_data(new.cond[i,1:4], size, and.logic,
paste0(row.names(new.cond)[i],'_sample_size_',size,'_simulated_data.csv'))
 i = 4
 nand.results <- simulate_data(new.cond[i,1:4], size, nand.logic,
paste0(row.names(new.cond)[i],'_sample_size_',size,'_simulated_data.csv'))
 i = 5
 xor.results <- simulate_data(new.cond[i,1:4], size, xor.logic,
paste0(row.names(new.cond)[i],'_sample_size_',size,'_simulated_data.csv'))
 i = 6
 xnor.results <- simulate_data(new.cond[i,1:4], size, xnor.logic,
paste0(row.names(new.cond)[i],'_sample_size_',size,'_simulated_data.csv'))
 i = 7
 imply.results <- simulate_data(new.cond[i,1:4], size, imply.logic,
paste0(row.names(new.cond)[i],'_sample_size_',size,'_simulated_data.csv'))
 i = 8
 nimply.results <- simulate_data(new.cond[i,1:4], size, nimply.logic,
paste0(row.names(new.cond)[i],'_sample_size_',size,'_simulated_data.csv'))
 i = 9
 or1.results <- simulate_data(new.cond[i,1:4], size, or.logic,
paste0(row.names(new.cond)[i],'_sample_size_',size,'_simulated_data.csv'))
}
```

```
##### STATISTICS ####

for (i in 1:length(row.names(new.cond))){ #
```

```
  fileName = paste0(row.names(new.cond)[i],'_sample_size_',size,'_simulated_data.csv')
  working.df <- read.csv(fileName, header=TRUE, row.names = 1)
  working.lmer <- lmer(Results ~ on + (1|group), data = working.df)

  cat(paste0('############ Reading filename ',fileName))
  print(confint(summary(glht(working.lmer,linfct=c('on==0')))))
  print(summary(glht(working.lmer,linfct=c('on==0'))))

}
```

```
##### MONTE CARLO SIMULATIONS #####

n.mc <- 1e4 # number of monte carlo simulations
n.to.try <- c(3,5,30)

beta.or <- matrix(nrow=length(n.to.try),ncol=n.mc)
beta.nor <- matrix(nrow=length(n.to.try),ncol=n.mc)
beta.and <- matrix(nrow=length(n.to.try),ncol=n.mc)
beta.nand <- matrix(nrow=length(n.to.try),ncol=n.mc)

beta.xor <- matrix(nrow=length(n.to.try),ncol=n.mc)
beta.xnor <- matrix(nrow=length(n.to.try),ncol=n.mc)
beta.imply <- matrix(nrow=length(n.to.try),ncol=n.mc)
beta.nimply <- matrix(nrow=length(n.to.try),ncol=n.mc)
```

```
# OR GATE

for (j in 1:length(n.to.try)){ # for each sample size
  n <- n.to.try[j]
  for (k in 1:n.mc){ # monte carlo simulation

    working.df <- simulate_data(new.cond[1,1:4],n,or.logic,"",write.file=FALSE)
    working.lmer <- lmer(Results ~ on + (1|group), data = working.df)
    beta.estimate <- (coef(working.lmer))$group$on1[1]

    beta.or[j,k] <- beta.estimate

  }
}
beta.or <- data.frame(beta.or,row.names = n.to.try)
colnames(beta.or) <- 1:n.mc
beta.or <- t(beta.or)

write.csv(beta.or,'montecarlo_betas_or_gate.csv')
```

```
# NOR GATE

for (j in 1:length(n.to.try)){ # for each sample size
  n <- n.to.try[j]
  for (k in 1:n.mc){ # monte carlo simulation

    working.df <- simulate_data(new.cond[2,1:4],n,nor.logic,"",write.file=FALSE)
    working.lmer <- lmer(Results ~ on + (1|group), data = working.df)
    beta.estimate <- (coef(working.lmer))$group$on1[1]

    beta.nor[j,k] <- beta.estimate

  }
}
beta.nor <- data.frame(beta.nor,row.names = n.to.try)
colnames(beta.nor) <- 1:n.mc
beta.nor <- t(beta.nor)

write.csv(beta.nor,'montecarlo_betas_nor_gate.csv')
```

```
# AND GATE

for (j in 1:length(n.to.try)){ # for each sample size
  n <- n.to.try[j]
  for (k in 1:n.mc){ # monte carlo simulation

    working.df <- simulate_data(new.cond[3,1:4],n,and.logic,"",write.file=FALSE)
    working.lmer <- lmer(Results ~ on + (1|group), data = working.df)
    beta.estimate <- (coef(working.lmer))$group$on1[1]

    beta.and[j,k] <- beta.estimate
  }
}
beta.and <- data.frame(beta.and,row.names = n.to.try)
colnames(beta.and) <- 1:n.mc
beta.and <- t(beta.and)

write.csv(beta.and,'montecarlo_betas_and_gate.csv')
```

```
# NAND GATE

for (j in 1:length(n.to.try)){ # for each sample size
  n <- n.to.try[j]
  for (k in 1:n.mc){ # monte carlo simulation

    working.df <- simulate_data(new.cond[4,1:4],n,c(1,0,0,0),"",write.file=FALSE)
    working.lmer <- lmer(Results ~ on + (1|group), data = working.df)
    beta.estimate <- (coef(working.lmer))$group$on1[1]

    beta.nand[j,k] <- beta.estimate
  }
}
beta.nand <- data.frame(beta.nand,row.names = n.to.try)
colnames(beta.nand) <- 1:n.mc
beta.nand <- t(beta.nand)

write.csv(beta.nand,'montecarlo_betas_nand_gate.csv')
```

```
# XOR GATE

beta <- matrix(nrow=length(n.to.try),ncol=n.mc)
fileName <- 'montecarlo_betas_xor_gate.csv'
gate_text <- 'XOR'
logic <- c(0,1,1,0)

for (j in 1:length(n.to.try)){ # for each sample size
  n <- n.to.try[j]
  for (k in 1:n.mc){ # monte carlo simulation

    working.df <- simulate_data(new.cond[gate_text,1:4],n,logic,"",write.file=FALSE)
    working.lmer <- lmer(Results ~ on + (1|group), data = working.df)
    beta.estimate <- (coef(working.lmer))$group$on1[1]

    beta[j,k] <- beta.estimate
  }
}
beta <- data.frame(beta,row.names = n.to.try)
colnames(beta) <- 1:n.mc
beta <- t(beta)
write.csv(beta,fileName)
```

```
# XNOR GATE

beta <- matrix(nrow=length(n.to.try),ncol=n.mc)
fileName <- 'montecarlo_betas_xnor_gate.csv'
```

```
gate_text <- 'XNOR'
logic <- c(1,0,0,1)

for (j in 1:length(n.to.try)){ # for each sample size
  n <- n.to.try[j]
  for (k in 1:n.mc){ # monte carlo simulation

    working.df <- simulate_data(new.cond[gate_text,1:4],n,logic,"",write.file=FALSE)
    working.lmer <- lmer(Results ~ on + (1|group), data = working.df)
    beta.estimate <- (coef(working.lmer))$group$on1[1]

    beta[j,k] <- beta.estimate
  }
}
beta <- data.frame(beta,row.names = n.to.try)
colnames(beta) <- 1:n.mc
beta <- t(beta)
write.csv(beta,fileName)
```

```
# IMPLY GATE

beta <- matrix(nrow=length(n.to.try),ncol=n.mc)
fileName <- 'montecarlo_betas_imply_gate.csv'
gate_text <- 'IMPLY'
logic <- c(1,0,1,1)

for (j in 1:length(n.to.try)){ # for each sample size
  n <- n.to.try[j]
  for (k in 1:n.mc){ # monte carlo simulation

    working.df <- simulate_data(new.cond[gate_text,1:4],n,logic,"",write.file=FALSE)
    working.lmer <- lmer(Results ~ on + (1|group), data = working.df)
    beta.estimate <- (coef(working.lmer))$group$on1[1]

    beta[j,k] <- beta.estimate
  }
}
beta <- data.frame(beta,row.names = n.to.try)
colnames(beta) <- 1:n.mc
beta <- t(beta)
write.csv(beta,fileName)
```

```
# NIMPLY GATE

beta <- matrix(nrow=length(n.to.try),ncol=n.mc)
fileName <- 'montecarlo_betas_nimply_gate.csv'
gate_text <- 'NIMPLY'
logic <- c(0,1,0,0)

for (j in 1:length(n.to.try)){ # for each sample size
  n <- n.to.try[j]
  for (k in 1:n.mc){ # monte carlo simulation

    working.df <- simulate_data(new.cond[gate_text,1:4],n,logic,"",write.file=FALSE)
    working.lmer <- lmer(Results ~ on + (1|group), data = working.df)
    beta.estimate <- (coef(working.lmer))$group$on1[1]

    beta[j,k] <- beta.estimate
  }
}
beta <- data.frame(beta,row.names = n.to.try)
colnames(beta) <- 1:n.mc
beta <- t(beta)
write.csv(beta,fileName)
```

**Table S10. Script for analyzing data in Figure 5.**

```
setwd('/Users/)

set.seed(2024)
library(lme4)
library(tidyr)
library(effects)
library(lmerTest)
library(multcomp)

REU.df <- read.csv("data2.csv",header=TRUE)
strength <- c('Trad','Med','Med_Low','Low','LuxR','Cons') # this is i
type <- c('Minus','aTc','OC6','Plus')

for(i in 1:length(strength)){

  working_strength <- strength[i]

  working_minus_label <- paste0(working_strength,'_',type[1])
  working_aTC_label <- paste0(working_strength,'_',type[2])
  working_OC6_label <- paste0(working_strength,'_',type[3])
  working_plus_label <-paste0(working_strength,'_',type[4])

  working_minus <- na.omit(REU.df[working_minus_label])
  working_aTC <- na.omit(REU.df[working_aTC_label])
  working_OC6 <- na.omit(REU.df[working_OC6_label])
  working_plus <- na.omit(REU.df[working_plus_label])

  Results <- c(t(working_minus), t(working_aTC), t(working_OC6), t(working_plus))
  group <-
as.factor(c(rep(1,dim(working_minus)[1]),rep(2,dim(working_aTC)[1]),rep(3,dim(working_OC6)[1]),rep(4,d
im(working_plus)[1])))
  on <- as.factor(c(rep(0,dim(working_minus)[1]),rep(1,(length(Results)-dim(working_minus)[1]))))

  working.df <- data.frame(Results,group,on)

  cat(paste("########### Now doing strength",working_strength,'\n'))

  working.lmer <- lmer(Results ~ on + (1|group), data = working.df)

  print(confint(summary(glht(working.lmer,linfct=c('on1==0')))))
  print(summary(glht(working.lmer,linfct=c('on1==0'))))

}
```

**Table S11. Monte Carlo Script for power analysis**

```
setwd('/Users/ ')
source("simulate_data_var.R") # allows to pass a variance input, fraction of the mean response.

set.seed(2024)

library('lme4')
library('effects')
library('broom')
library('multcomp')

k.means <- read.csv('20240806_data_k_6.csv',row.names=1,header=TRUE)
or.logic <- c(0, 1, 1, 1)

n.mc <- 1e4 # number of monte carlo simulations
n.to.try <- 2:30
group_numbers <- c('0','1','2','3','4','5')

variance.values <- c(0.1, 0.2, 0.3, 0.4, 0.5, 0.6, 0.7, 0.8, 0.9, 1)

for (i in 1:length(group_numbers)){

  for (m in 1:length(variance.values)){ # for each variance

    group_number <- group_numbers[i]
    beta.store <- matrix(nrow=length(n.to.try),ncol=n.mc) # init
    p.value <- matrix(nrow=length(n.to.try),ncol=n.mc) # init
    variance <- variance.values[m]

    for (j in 1:length(n.to.try)){ # for each sample size

      n <- n.to.try[j]
      for (k in 1:n.mc){ # monte carlo simulation

        working.df <-
simulate_data_var(k.means[group_number,1:4],n,or.logic,"",variance,write.file=FALSE,output.text=FALSE)
        working.lmer <- lmer(Results ~ on + (1|group), data = working.df)

        beta.estimate <- (coef(working.lmer))$group$on1[1]
        beta.store[j,k] <- beta.estimate

        answer <- summary(glht(working.lmer,linfct=c('on1==0')))
        p.value[j,k] <- answer$test$pvalues[1]
      }
      cat(paste0('Done with sample size ',n,' in gate number ',group_number,' and variance fraction
',variance,'.\n'))
    }

    beta.store <- data.frame(beta.store,row.names = n.to.try)
    colnames(beta.store) <- 1:n.mc
    beta.store <- t(beta.store)

write.csv(beta.store,paste0('montecarlo_betas_kmeans_group_',group_number,'variance_frac_',variance,'.
csv'))
    p.value <- data.frame(p.value,row.names = n.to.try)
    colnames(p.value) <- 1:n.mc
    p.value <- t(p.value)

write.csv(p.value,paste0('montecarlo_p_values_kmeans_group_',group_number,'variance_frac_',variance,'
.csv'))
    }
}
```

**Supplementary Table S12.** Benchling files and plasmid maps for constructs from Figure 5.

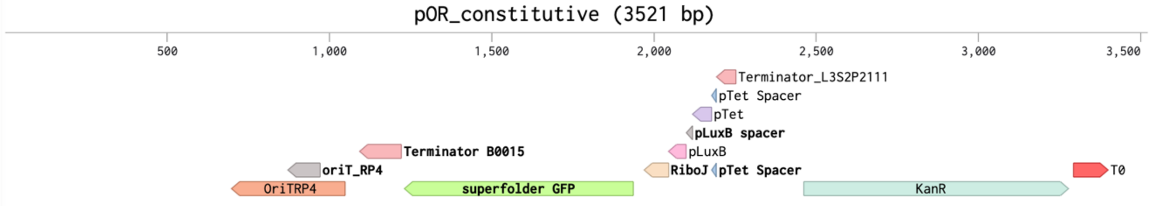


https://benchling.com/s/seq-axLTXACiq0rWTqYHDlq7?m=slm-bm5kERWa6BkQVY0aIfNE

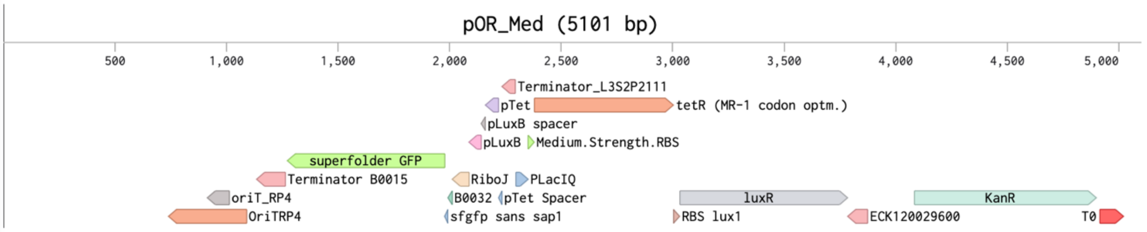


https://benchling.com/s/seq-dZ6LKBDeak1FYVYaG0iS?m=slm-s8XmtRiiwGIDfHmqlF78

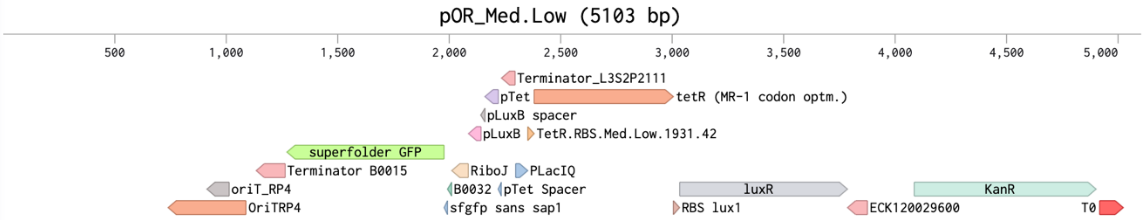


https://benchling.com/s/seq-aAK0RNRyoEXXgMu2PgtZ?m=slm-n7BeNTVr3ub03lQPGfce

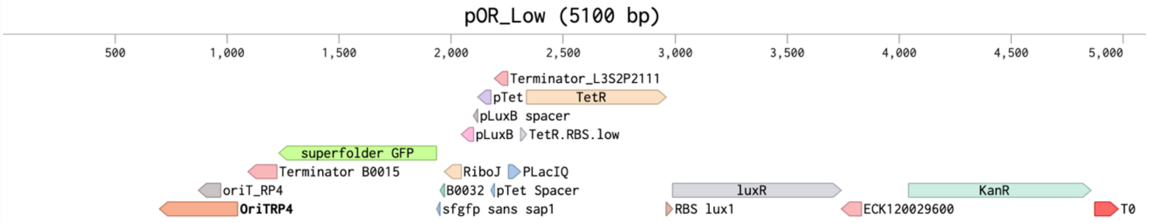


https://benchling.com/s/seq-jx7FbPk2kKXjOLmrs3NT?m=slm-mCRndI4unYUdpxA5d1oJ

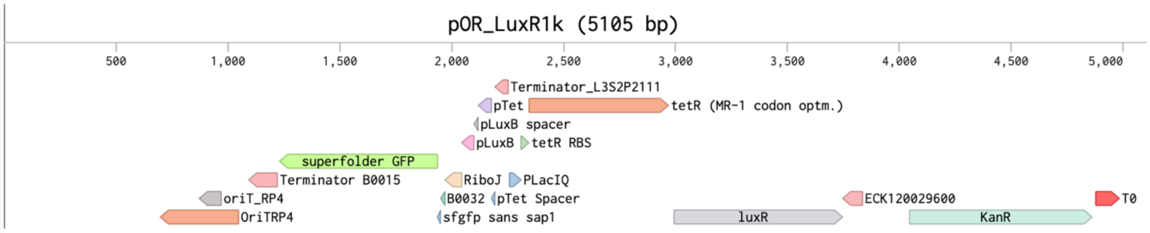


https://benchling.com/s/seq-HvPFUAQ6q3jM8ILJpmLj?m=slm-tSloQeRd08Pmr3ieE3zU

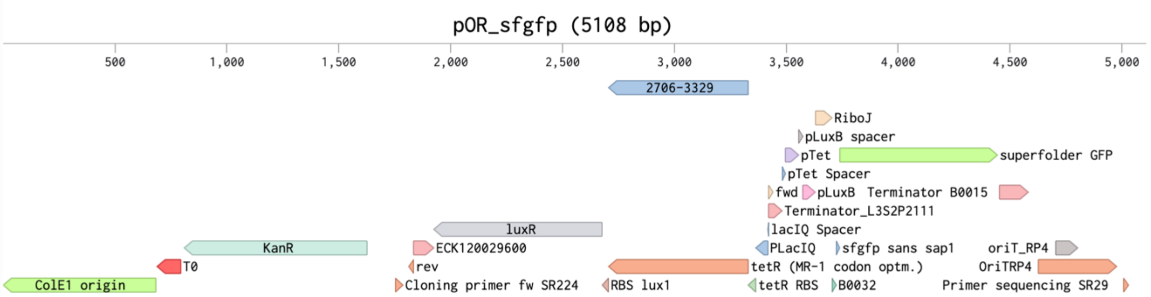


https://benchling.com/s/seq-4ufeLgf8lcAqqnppexFh?m=slm-Ld7T8gF1I3OrOXGbIvbs
